# Th1 polarization in *Bordetella pertussis* vaccine responses is maintained through a positive feedback loop

**DOI:** 10.1101/2024.08.05.606623

**Authors:** Lisa Willemsen, Jiyeun Lee, Pramod Shinde, Ferran Soldevila, Minori Aoki, Shelby Orfield, Mari Kojima, Ricardo da Silva Antunes, Alessandro Sette, Bjoern Peters

**Affiliations:** Center for Infectious Disease and Vaccine Research, La Jolla Institute for Immunology, La Jolla, CA, USA; Department of Medicine, University of California San Diego, La Jolla, CA, USA

**Keywords:** Vaccine, Bordetella pertussis, immune response, T cells, Th1 polarization

## Abstract

Outbreaks of *Bordetella pertussis* (BP), the causative agent of whooping cough, continue despite broad vaccination coverage and have been increasing since vaccination switched from whole-BP (wP) to acellular BP (aP) vaccines. wP vaccination has been associated with more durable protective immunity and an induced Th1 polarized memory T cell response. Here, a multi-omics approach was applied to profile the immune response of 30 wP and 31 aP-primed individuals and identify correlates of T cell polarization before and after Tdap booster vaccination. We found that transcriptional changes indicating an interferon response on day 1 post-booster along with elevated plasma concentrations of IFN-γ and interferon-induced chemokines that peaked at day 1-3 post-booster correlated best with the Th1 polarization of the vaccine-induced memory T cell response on day 28. Our studies suggest that wP-primed individuals maintain their Th1 polarization through this early memory interferon response. This suggests that stimulating the interferon pathway during vaccination could be an effective strategy to elicit a predominant Th1 response in aP-primed individuals that protects better against infection.

## Introduction

Whooping cough remains a significant and worldwide public health concern marked by periodic outbreaks and regularly life-threatening complications in young infants(1, 2). Whooping cough is caused by a respiratory infection with the *Bordetella pertussis* (BP) bacteria. In 1950, the initial vaccine, the alum-adjuvanted ‘whole cell’ (wP) inactivated BP vaccine, was administered and significantly reduced the disease incidence. However, due to adverse effects, the wP vaccine was replaced by an acellular pertussis (aP) vaccine. The aP vaccine contains different BP antigens including inactivated pertussis toxin (PT) and cell surface proteins of BP including filamentous hemagglutinin (FHA), fimbriae 2/3 (Fim2/3), and pertactin (PRN) and is administered together with tetanus toxoid (TT) and diphtheria toxoid (DT) as alum-adjuvanted DTaP vaccine to infants and as Tdap vaccine to teenagers and adults(3, 4). While the aP vaccine is generally associated with fewer adverse effects, the wP vaccine induces a broader and more robust immune response that lasts longer(5, 6). Interestingly, receipt of at least 1 wP dose significantly improves the durability of immunity despite subsequent aP doses as compared to exclusively aP priming and boosting(6). Although both the aP and wP vaccines protect against disease, the wP vaccine protects significantly better against infection than the aP vaccine(7, 8). The current resurgence of whooping cough has been linked to the switch to aP-priming(9-13). This resurgence may be explained by the distinct immune profiles elicited by these vaccines.

CD4^+^ T cell immunity is required to protect against BP infection(3, 14). However, the type of T helper response between aP and wP-primed individuals differs and is each characterized by specific cytokine profiles and effector functions that coordinate immune responses to BP. Compared to wP vaccine priming, aP vaccine priming elicits a more Th2-polarized immune response(15-19) which is recognized by the production of interleukin-4 (IL-4), IL-5, IL-10, and IL-13 and associated with a humoral response including the production of IgE antibodies(20). Furthermore, Th2-polarized cells ensure the recruitment and activation of eosinophils and mast cells, thereby contributing to allergic inflammation(21). Compared to aP-primed individuals, wP-primed individuals show an increased Th1 over Th2-polarized immune response against BP which persists in adulthood despite aP booster vaccination(15-19). Th1-polarized cells primarily produce interferon-gamma (IFN-γ), tumor necrosis factor (TNF), and IL-2 and elicit both antibody-mediated and cell-mediated immunity(22). Th1 cells promote the phagocytosis and killing of pathogens by macrophages, enhance antigen presentation by dendritic cells, and stimulate the differentiation of cytotoxic T cells. Using a mouse model, research has shown that optimum protection against BP requires induction of a Th1, but not a Th2-polarized response(23). Additionally, natural infection with BP in children also induced a Th1-biased response(24).

T cell polarization starts with antigen-presenting cells (APC) presenting antigens via HLA-II to naive or memory T cells with a matching T cell receptor. This process can take place either at the local site of antigen exposure or within the nearby draining lymph nodes. After antigen recognition and co-stimulation, APC secrete different cytokines which are the first triggers of T cell polarization. Researchers have described that IL-12(25-28) and IL-18(29) are important Th1 polarizing cytokines, while IL-4 is known to stimulate Th2 polarization(30). The Th1 polarizing cytokines IL-12 and IL18 initiate the production of IFN-γ by natural killer (NK) and CD8^+^ T cells within 2-6 hours after antigen exposure(31, 32). This early derived IFN-γ is essential for controlling certain infections before the remaining IFN-γ gets produced by other T cells. Early IFN-γ directly activates macrophages to enhance their capacity to eliminate pathogens but also regulates the differentiation of CD4^+^ T cells into Th1 cells that will also start producing IFN-γ(33). Additionally, after booster vaccination or reinfection, IFN-γ will also be quickly produced by reactivated memory CD4^+^ T cells which also contributes to Th1 polarization of naive T cells(34, 35).

Previous research showed that BP infection is more severe in IFN-γ-depleted mice(36). Other researchers found that NK cell depletion caused disseminated lethal lung infection in BP-infected mice(37). This severe outcome was associated with a reduced antigen-specific Th1 response and increased Th2 response. Interestingly, Gillard et al. found that transcriptional antiviral responses on day 1 post-vaccination with Tdap-IPV (inactivated Poliovirus) were positively associated with antibody responses up to 1 year post-vaccination (38).

The mechanism of how wP-primed individuals maintain their Th1 polarization despite repeated aP boosters remains a puzzle(39). To identify factors contributing to increased Th1 polarization, we applied a multi-omics approach to profile the immune response of 31 aP and 30 wP-primed individuals longitudinally in the first 2 weeks post-vaccination and identified correlates of T cell polarization before and after Tdap booster vaccination. We confirmed the increased Th1 polarization in the wP individuals of our cohort and found that Th1 polarization pre-vaccination correlated positively with the vaccine-induced interferon response in plasma and PBMCs on day 1-3 post-booster on transcriptional and protein levels, which in turn correlated with a maintained Th1 polarization post-booster response. This suggested that changing the BP vaccination strategy by targeting the interferon pathway might boost Th1 responses against BP which could improve protection against infection and vaccine durability.

## Results

### Longitudinal samples from an adult cohort were collected to study Tdap booster vaccination-induced immune responses

We recruited healthy adults primed with either the aP (n=31) or wP (n=30) vaccine during infancy (**Fig. 1A**). Study participants did not receive Tdap booster vaccinations four years prior to study enrollment and aP and wP groups were matched by sex (**Supplementary Table 1**). From these participants, longitudinal blood samples were taken before (days - 31, -14, and 0) and after Tdap booster vaccination (days 1, 3, 7, 14, and 28; **Fig. 1B**). The multiple pre-vaccination samples were utilized to establish robust baseline measurements per donor and were used to calculate post-booster responses. Previously, a multi-omics approach was applied to evaluate the immune response to BP vaccination by integrating plasma antigen-specific IgG levels including IgG isotypes (IgG1-4), plasma concentrations of 45 different cytokines, and peripheral blood mononuclear cell (PBMC) subset frequencies and transcriptomics(40). Here we added measurements of T cell activation and polarization and subsequent analysis of the same donors at multiple time points (**Fig. 1B, Supplementary Table 2**). The increased plasma levels of IgGs specifically targeting the Tdap vaccine antigens PT, PRN, FHA, FIM2/3, TT, and DT on day 7 and 14 post-booster vaccination verified an induction of a humoral response and confirmed successful booster immunization ( **Supplementary Fig. 1A**).

**Figure 1.**
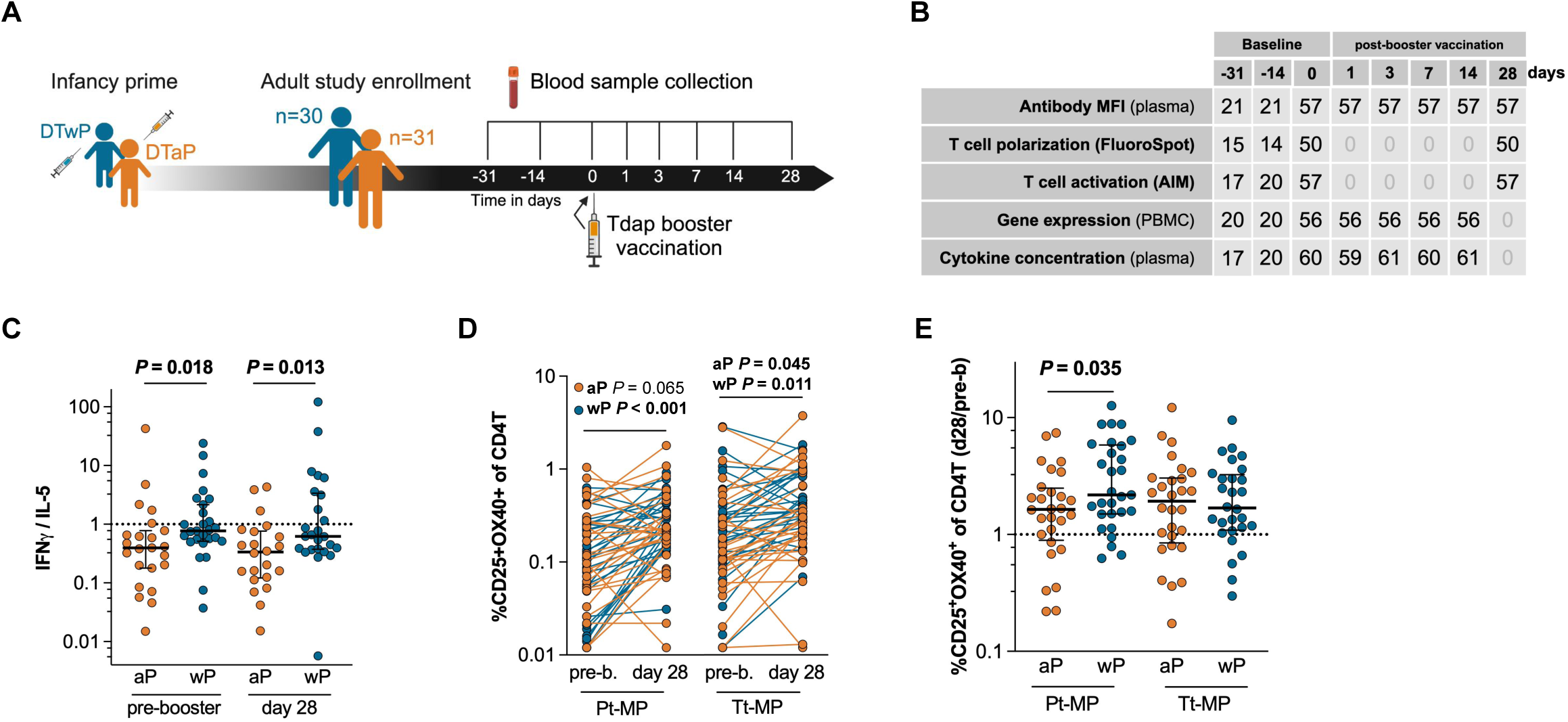
Whole-cell pertussis vaccine priming increases Th1 polarization despite acellular booster vaccination. **A)** Schematic illustration of Tdap booster vaccination study outline. Healthy adults were recruited who were primed with either the DTaP (n=31) or DTwP (n=30) vaccine during infancy. Blood was collected before and after Tdap booster vaccination. **B**) Table indicating sample quantity per experiment type and time point. Plasma was used for antibody and cytokine measurements and peripheral blood mononuclear cells (PBMCs) were used to determine gene expression, and T cell activation and polarization. **C**) IFN-γ and IL-5 producing cells (spot-forming cells, SFC) were measured by Fluorospot after cells were stimulated with aP vaccine antigens for 14 days and derived from blood sampled before and 28 days after Tdap booster vaccination (black lines represent the median with interquartile range). The IFN-γ/IL-5 ratio was used as a measurement of Th1/Th2 polarization. (n=22-23 aP, 25-27 wP, P-values were calculated by multiple two-tailed Mann-Whitney tests). **D-E**) An activation induced marker (AIM) assay was performed in which PBMCs were stimulated with 1 μg/mL of either the pertussis or tetanus peptide megapools for 18-24 hours. CD25^+^ and OX40^+^ CD4^+^ T cells were measured by flow cytometry and shown as percentage of total CD4^+^ T cells (**D**) and as 28 days post/pre booster vaccination ratio (**E**) where black lines represent the median with interquartile range. n=28 aP, 29 wP, P-values were calculated by **D**) multiple two-tailed Wilcoxon matched-pairs test and **E**) multiple two-tailed Mann-Whitney tests). **C-E**) Shapiro-Wilk tests were performed to assess normality.

### Whole-cell pertussis vaccine priming increases Th1 polarization despite acellular booster vaccination

Previous studies have shown that priming with the wP vaccine in infancy is associated with a life-long increase in Th1 polarization of the vaccine response compared to aP-primed individuals that show an increased Th2 polarization(15-19). We evaluated the Th1/Th2 polarization ratio in our cohort using a FluoroSpot assay(16) quantifying cytokine-secreting cells responding to aP vaccine antigen(15) stimulation following a 14-day expansion period. We selected IFN-γ and IL-5 as representative cytokines for Th1 and Th2 responses respectively(16). The Th1/Th2 (IFN-γ/IL-5) polarization ratio, further referred to as Th1 polarization, was significantly increased in wP compared to aP-primed participants before and 28 days after Tdap booster vaccination (**Fig. 1C, Supplementary Fig. 1B**) and these differences were independent of age after booster vaccination ( **Supplementary Fig. 1C**). Next, to identify vaccine-responsive CD4^+^ T cells, the co-expression of the T cell activation surface markers interleukin 2 receptor alpha (CD25) and TNF Receptor Superfamily Member 4 (OX40) was determined of CD4^+^ T cells before and 28 days post-booster vaccination by applying an activation induced marker (AIM) assay(41). In wP-primed participants, the percentage of activated pertussis and tetanus antigen-specific CD4+ T cells was significantly increased post-vs pre-booster vaccination (**Fig. 1D**). aP-primed individuals’ tetanus antigen-specific CD4+ T cell response was significantly increased and their pertussis antigen-specific CD4+ T cell response was close to the significance threshold (P=0.065). Interestingly, the AIM+ CD4^+^ T cell response against pertussis antigens of wP-primed participants showed a significantly greater increase compared to aP-primed participants, while no differences were observed regarding the AIM+ CD4+ T cell response boost against tetanus antigens (**Fig. 1E**). The greater increased anti-pertussis T cell response in wP participants mainly originated from the stronger post-booster response in wP participants (**Supplementary Fig. 1D**). To conclude, the previously described cell-mediated immunity against aP antigens and the Th1 polarization shift(15-19) in wP-primed individuals were confirmed in the wP-primed participants of this cohort.

### Interferon signaling day 1 post-booster vaccination positively correlates with Th1 polarization

To create a comprehensive picture of the immune response induced by Tdap booster vaccination, RNAseq was performed on PBMCs before and 1, 3, 7, and 14 days after booster vaccination. aP and wP-primed individuals were pooled for this analysis. RNAseq analysis identified a combined set of 1938 differentially expressed genes (DEGs) that showed significantly different expression at one or more of the post-booster time points compared to baseline (**Fig. 2A**, FDR<0.05). The highest number of DEGs was recorded for day 1, with 579 downregulated and 922 upregulated genes that accounted together for 75.5% of the DEGs. In contrast, only 0.3% of DEGs were exclusive to day 3, 22.7% for day 7, and 1.6% for day 14. The most significant DEGs on day 1 included the increased expression of interferon-associated transcripts including *FCGR1A, STAT1, IRF1, GBP1,* and *CXCL10.* Day 7 DEGs included the increased expression of transcripts encoding immunoglobulin segments, such as *IGHG1, IGHV6-1*, *IGHGP*, *IGHG4*, and *IGKC,* confirming effective humoral booster vaccination, now on the RNA level.

**Figure 2.**
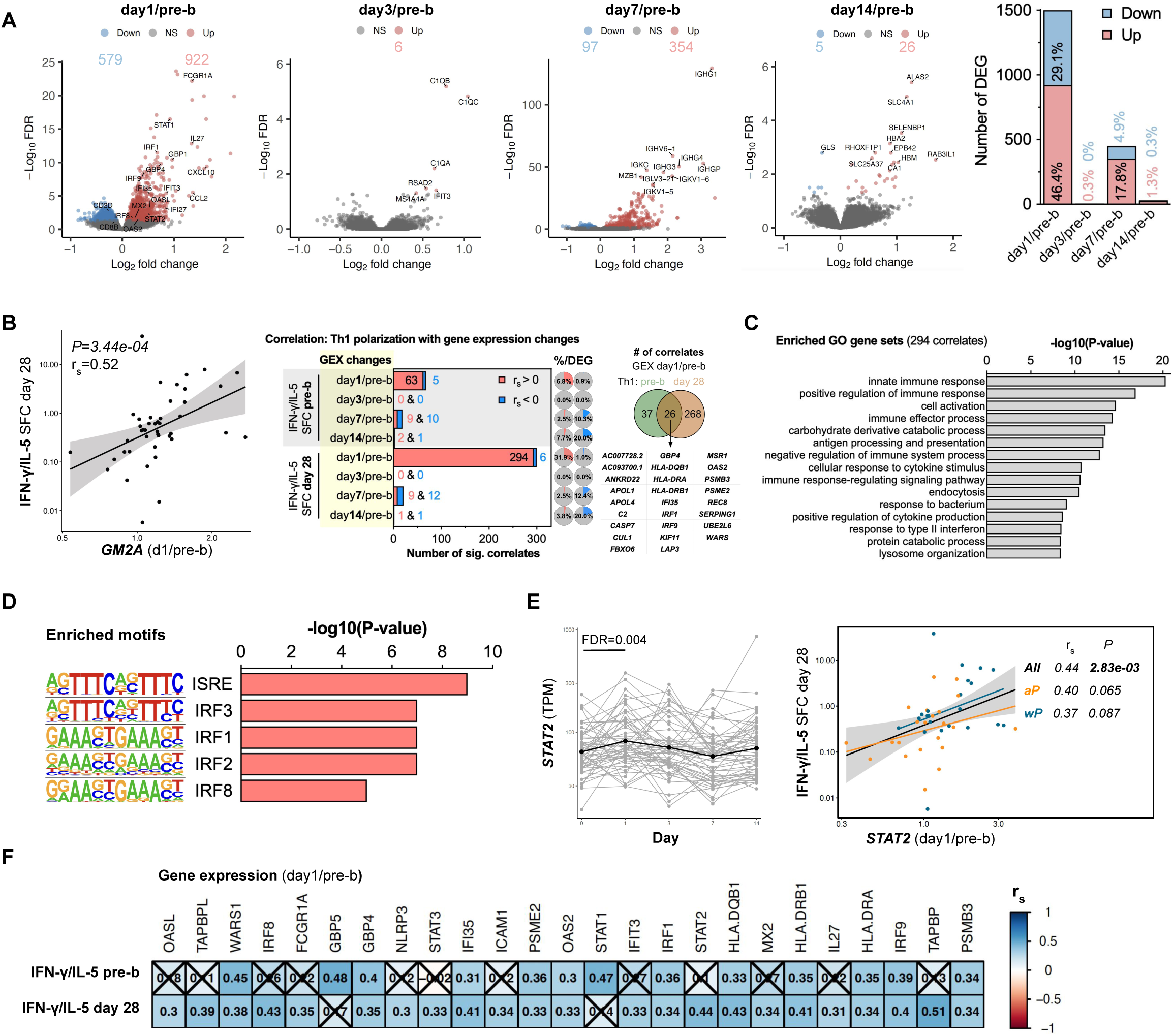
Booster-induced interferon signaling positively correlates with Th1 polarization. **A)** Volcano plots showing the log_2_(fold-change) and -log_10_(FDR) of the differentially expressed genes (DEG) of day 1, 3, 7, 14 post/pre-booster vaccination with downregulated genes in blue and upregulated genes in red (FDR<0.05). The number and percentage of DEG of the total analysis are depicted in the bar chart. **B**) Spearman sample correlation analysis was performed using the DEG changes and T cell polarization (IFN-γ/IL-5 SFC). A dotplot of most significant correlate (*GM2A* day 1/pre-booster with Th1 polarization on day 28 post-booster) is depicted and the number of significant correlates per time point comparison (*P*<0.05) and its percentage of the total number of DEG of that time point. On the right is a venn diagram depicting the number of correlates from the DEG changes on day1/pre-b that enclosed a significant correlation with Th1 polarization pre vs post-booster including a list of the overlapping correlates. **C)** The top 15 gene sets identified by Gene Ontology (GO) gene set enrichment analysis using the 294 DEG (day 1/pre-booster) that also showed a positive correlation with the T cell polarization on day 28. **D**) The top 5 known motifs identified by HOMER motif enrichment analysis using the 294 DEG (day 1/pre-booster) that also showed a positive correlation with the T cell polarization on day 28. **E**) *STAT2* gene expression (TPM) over time with the black line indicating the mean expression (left) and Spearman sample correlation (right) between *STAT2* (day 1/pre-booster) and T cell polarization data 28 days post booster (IFN-γ/IL-5 SFC). Spearman dot plot was created with aP and wP groups combined (“All”) and separately. **F**) Heatmap depicting c of selected transcriptional changes (day1/pre-booster) vs T cell polarization data pre- and post-booster. Insignificant correlates are crossed (*P*>0.05). RNA n=56, correlation analysis n=44. **A-F**) aP and wP-primed individuals were pooled for this analysis.

Next, Spearman correlation analysis was performed to identify which of the DEGs were associated with Th1 polarization. aP and wP-primed individuals were pooled for this analysis since the aim was to find correlates of a Th1 polarized anti-BP T cell response and not to further identify differences between aP and wP-primed individuals. Moreover, additional factors, other than infancy priming, may affect T cell polarization, such as an individual’s history of asymptomatic infections or undiagnosed colonization or symptomatic infections. This correlation analysis identified, for example, that transcriptional changes on day 1 vs pre-booster (day 1/pre-b) in Ganglioside GM2 Activator (*GM2A*), encoding a small glycolipid transport protein, were highly correlated with Th1 polarization on day 28 (**Fig. 2B, left panel**). By performing these analyses systematically, we found that most transcriptional changes that showed a significant correlation (*P*<0.05) with Th1 polarization originated from day 1/pre-b DEG together with Th1 polarization 28 days post-booster, both absolutely and relatively to its number of DEG (**Fig. 2B, middle panel**). Almost all (294/300) day 1/pre-b transcriptional perturbations that correlated with Th1 polarization 28 days post-booster enclosed a positive association. Although the majority of Th1 correlates included Th1 polarization 28 days post-booster, significant associations including Th1 polarization pre-booster also mainly originated from day 1/pre-b DEG and were mostly positive correlations (63/68). Of these 63 positive correlates, 26 were also significantly associated with Th1 polarization 28 days post-booster (**Fig. 2B, right panel**). All significant Th1-DEG correlates (*P*<0.05) are listed in **Supplementary Table 3**.

The 294 DEGs identified on day 1 with a positive correlation with Th1 polarization 28 days post-booster were examined for shared properties that explain their coordinated upregulation. Gene set enrichment analysis (GSEA) using Gene Ontology (GO) terms identified different immunological gene sets including innate immune response (GO:0045087), cell activation (GO:0001775, which includes T cell activation (GO:0042110)), immune effector process (adaptive immune response, GO:0002252) and response to type II interferon (GO:0034341, **Fig. 2C)**. Additionally, GSEA using the Reactome gene sets identified interferon signaling (R-HSA-913531) and antiviral mechanism by IFN-stimulated genes (R-HSA-1169410) among the top 15 most significantly enriched gene sets (**Fig. S2A**). Pathway enrichment analysis using the Kyoto Encyclopedia of Genes and Genomes (KEGG) pathways identified the pertussis (hsa05133) pathway as the third most significantly enriched (**Fig. S2C**). Separately, DNA motif enrichment analysis using the 294 DEGs identified interferon-stimulated response element (ISRE) and interferon regulatory factors (IRFs) as the most enriched motifs (**Fig. 2D**). This is corroborated by the gene expression of multiple interferon-related transcription factors including *STAT2, IRF1, IRF8, and IRF9*, all of which peak at day 1 post-booster and are correlated with increased Th1 polarization 28 days post-booster (**Fig. 2E-F, Supplementary Fig. 2C**). Correlation analysis was also performed with aP and wP separately. Positive correlation coefficients were found for IFN-related correlates in both groups. Although not all correlations were significant within the aP and wP groups, this analysis is likely underpowered yet suggests that our findings are not dominated by aP vs wP infancy priming. Other interferon-related Th1 polarization 28 days post-booster correlates are *FCGR1A, FI35, IFIT3, IRF8, IL27, OAS2, OASL, GBP4, and MX2* (**Fig. 2F**) and were induced on day 1 post-booster (**Fig. 2A**). Similarly, Th1 polarization pre-booster also correlated with changes in booster-induced interferon-related transcripts including GBP4, IFI35, OAS2, STAT1, IRF1, and IRF9. Taken together, Th1 polarization of BP-specific CD4^+^ T cells pre- and day 28 post-booster correlated strongly with the changes in interferon signaling detected in the transcriptional profile of PBMCs on day 1 post-booster vaccination.

### Plasma IFN-**γ** concentrations increase on day 1 post-booster vaccination and correlate with Th1 polarization

Next, we examined which vaccine-induced plasma cytokines correlated with Th1 polarization pre- and 28 days post-booster. The concentrations of 45 different cytokines were measured before and 1, 3, 7, and 14 days after booster vaccination to characterize the immune response induced by Tdap booster vaccination at the plasma protein level (**Fig. 3A)**. Of all cytokines and time points, the concentration of IFN-γ at day 1 post-booster showed the highest and most significant induction after booster vaccination (**Fig. 3A-B**). Fold changes of post/pre-booster were calculated per cytokine and post-booster time point and used for correlation analysis with the Th1 polarization data. Changes in plasma IFN-γ on day 1/pre-b significantly correlated with the Th1 polarization ratio both pre- and 28 days post-booster (**Fig. 3C, Supplementary Fig. 3A**, significant correlates (*P*<0.05) are listed in **Supplementary Table 4.**). While plasma IFN-γ levels peaked at day 1 post-booster, *IFNG* gene expression was not significantly increased on day 1 post-booster or any other time point measured (**Fig. 3D**). The presence of high levels of plasma IFN-γ on day 1 post-booster together with the lack of a transcriptional *IFNG* peak in PBMCs suggests that *IFNG* transcription occurred within hours post-booster vaccination and/or that IFN-γ is produced by tissue-resident cells present at the injection site or its draining lymph nodes. To conclude, plasma IFN-γ concentrations peak on day 1 post-booster vaccination and are positively associated with increased Th1 polarization pre- and 28 days post-booster.

**Figure 3.**
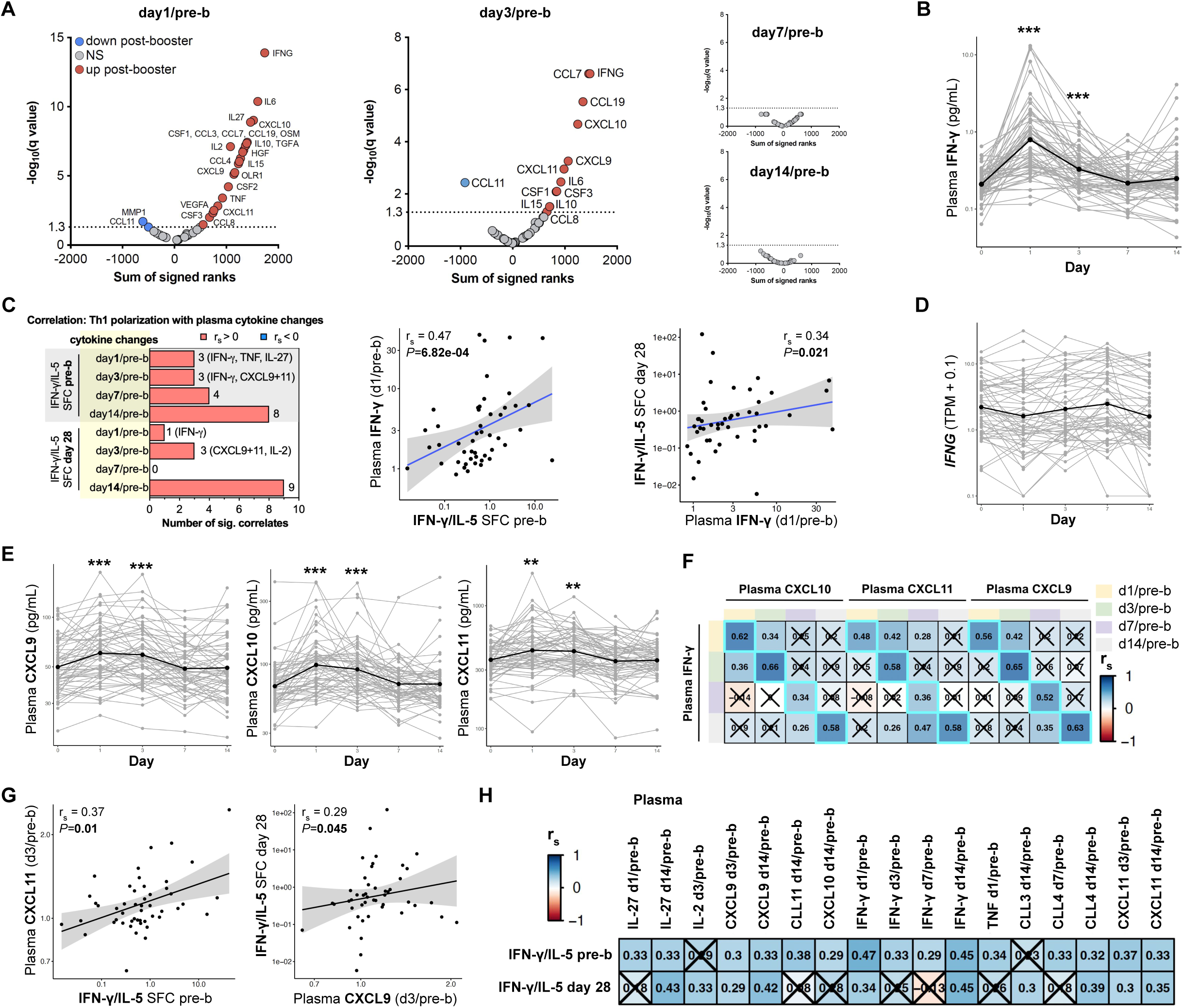
Plasma IFN-γ and its initiated chemokines increase post-booster and its changes positively correlate with Th1 polarization. **A)** Volcano plots showing the plasma cytokine concentration changes of day 1, 3, 7, 14 post/pre-booster vaccination with the sum of signed ranks of the two-tailed Wicoxon matched-paired signed rank test (x-axis) and -log_10_(q-value) on the y-axis. Cytokines that are significantly increased and decreased are highlighted in red and blue respectively (q-value<0.05). **B**) Plasma concentrations of IFN-γ (pg/mL) over time. **C)** Number of significant Spearman sample correlations (*P*<0.05) of plasma cytokine changes vs T cell polarization pre and 28 days post booster (IFN-γ/IL-5 SFC) and correlation dotplots of IFN-γ (day1/pre-b) vs T cell polarization pre and 28 days post booster (IFN-γ/IL-5 SFC). Spearman dot plots were created with aP and wP groups combined (“All”) and separately. **D)** IFNG gene expression (TPM +0.1) over time with the black line indicating the mean expression. **E**) Plasma concentrations of CXCL9, CXCL10, and CXCL11 over time. **F**) Heatmap depicting Spearman correlation coefficients (r_s_) of selected plasma cytokine changes showing that the dynamics of CXCL9-11 change with that of IFN-γ Insignificant correlates are crossed (*P*>0.05). **G**) *S*pearman sample correlation between either plasma CXCL9 (day3/pre-b) or CXCL11 (day3/pre-b) and T cell polarization data 28 days post booster (IFN-γ/IL-5 SFC). Spearman dot plots were created with aP and wP groups combined (“All”) and separately. **H**) Heatmap depicting Spearman correlation coefficients (r_s_) of selected plasma cytokine changes post/pre-b booster vaccination (columns) vs T cell polarization data before and after booster (rows). Insignificant results are crossed (*P*>0.05). Plasma cytokine n=59-61, RNA n=56, correlation analysis n=38-47. ***q-value<0.001, **q-value<0.01. **A-H**) aP and wP-primed individuals were pooled for this analysis.

### IFN-**γ** initiated chemokine production on days 1 and 3 positively associated with Th1 polarization

As changes in plasma IFN-γ and its transcriptional signaling on day 1 post-vaccination is the strongest correlate of Th1 polarization, we wanted to examine the behavior of other players in the IFN-γ pathway. IFN-γ is known to bind to the IFN-γ receptor which consists of the two subunits IFNGR1 and IFNGR2(42). When IFN-γ activates its receptor, downstream signaling ensures the phosphorylation of STAT1, which then forms a homodimer. These complexes translocate to the nucleus where they initiate transcription by binding to genes containing GAS elements in their promotors. CXCL9, CXCL10, and CXCL11 are examples of transcripts that will be transcribed after IFN-γ stimulation(43). These chemokines promote the migration of CXCR3-expressing cells, such as activated T cells and NK cells.

Interestingly, plasma levels of CXCL9, CXCL10, and CXCL11 significantly increased on day 1 and day 3 post-booster compared to pre-booster (**Fig. 3E**). Additionally, participants showing induction of plasma IFN-γ also showed a significant induction of CXCL9, CXCL10, and CXCL11, at any time point measured (**Fig. 3F**). The day 3/pre-b plasma concentrations of CXCL9 and CXCL11 positively correlated with Th1 polarization pre- and day 28 post-booster (**Fig. 3G-H, Supplementary Fig. 3B**). To conclude, we found evidence of a positive correlation between both systemic IFN-γ perturbations and interferon signaling with Th1 polarization at the protein level, mirroring the patterns observed in RNA transcripts.

### Tdap booster-induced changes in plasma IFN-**γ** and CXCL9-11 correlate with the magnitude of transcriptional IFN response

Since both changes in plasma IFN-γ and IFN-induced gene expression in PBMCs correlate with the Th1 polarization, we next investigated the relationship between the production of plasma IFN-γ and IFN-induced gene expression in PBMCs on day 1 post-booster vaccination. The production of plasma IFN-γ, CXCL9, CXCL10, and CXCL11 all significantly correlated with the gene expression magnitude of more than 20 IFN-associated genes in PBMCs (**Fig. 4A-B**). The 5 strongest correlations all involved plasma IFN-γ and significantly correlated with transcriptional changes in *STAT1, FCGR1A, GBP1, GBP2, and IRF1*. Moreover, changes in the IFN-induced plasma chemokines CXCL9, CXCL10, and CXCL11 also significantly correlated with these transcriptional changes.

**Figure 4.**
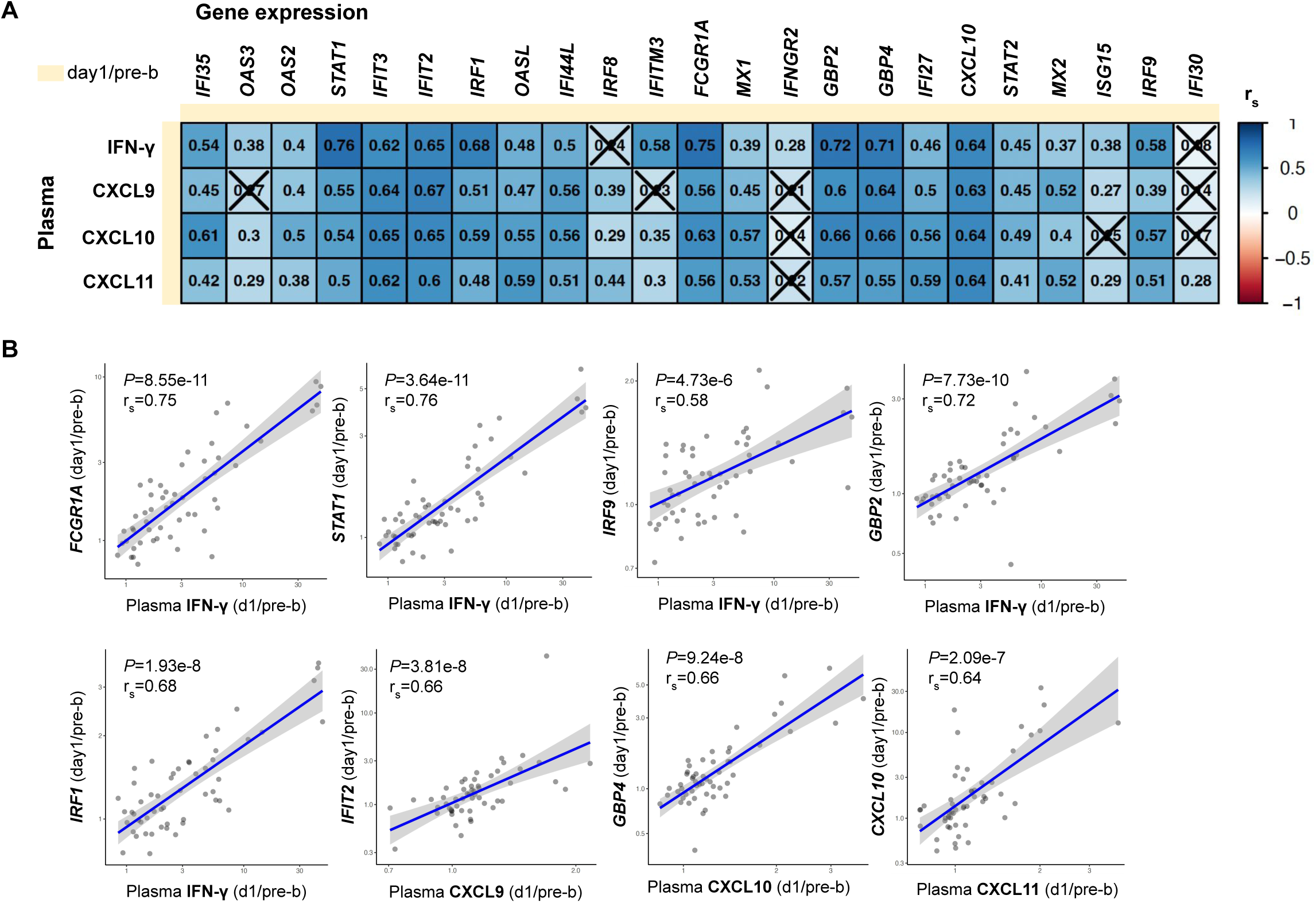
Changes in plasma IFN-γ and its initiated chemokines correlate with the magnitude of transcriptional IFN response in PBMCs. **A)** Heatmap depicting Spearman sample correlations coefficients (r_s_) of selected plasma cytokine changes with vaccine-induced genes d1/pre-b (DEG). Insignificant correlates are crossed (*P*>0.05). **B**) *S*pearman sample correlation between either plasma IFN-γ, CXCL9-11 (day1/pre-b) and selected IFN and vaccine-induced genes d1/pre-b (DEG). **A-B**) aP and wP-primed individuals were pooled for this analysis.

This confirms that individuals with increased plasma IFN-γ levels post-Tdap vaccination, also showed an increase in IFN-induced gene expression in PBMCs. These findings suggest that the vaccine-induced production of IFN-γ induced a systemic immune response in which IFN-associated gene expression was likely also initiated in non-T cell subsets. This subsequent systemic effect–which also correlated with the Th1 polarization– may contribute to maintaining the Th polarization differences observed in aP compared to wP individuals.

## Discussion

It is now well established that individuals primed during infancy with the wP vaccine maintain an increased Th1 polarization in their memory T cell response compared to aP-primed individuals (15-19). It has however remained a puzzle how this polarization is maintained over decades in wP-primed individuals despite repeated Tdap booster vaccinations. To address this question, we performed a longitudinal multi-omics analysis of the immune state pre- and post-Tdap booster vaccination to identify the cascade of events that maintains a Th1 polarized memory T cell response, which is associated with durable protection. In our hypothesis-generating study design, we comprehensively mapped the immune state by measuring many blood parameters, identified which ones were perturbed by booster vaccination, and then queried which changes were positively correlated with the Th1 response before and 28 days post-booster vaccination.

We verified the increased Th1 polarization in the wP individuals and showed that Th1 polarization pre-booster correlated positively with the booster-induced interferon response in PBMCs on day 1-3 post-booster on RNA and protein levels, which in turn positively correlated with the maintained Th1 polarization 28 days post-booster. This suggests that early IFN-γ secretion maintains the Th1 polarized response in wP-primed individuals despite multiple Tdap boosters, through a positive feedback loop (**Fig. 5**). Therefore, stimulating IFN-γ signaling during vaccination might boost Th1 responses against BP which could improve protection against infection and increase vaccine durability.

**Figure 5.**
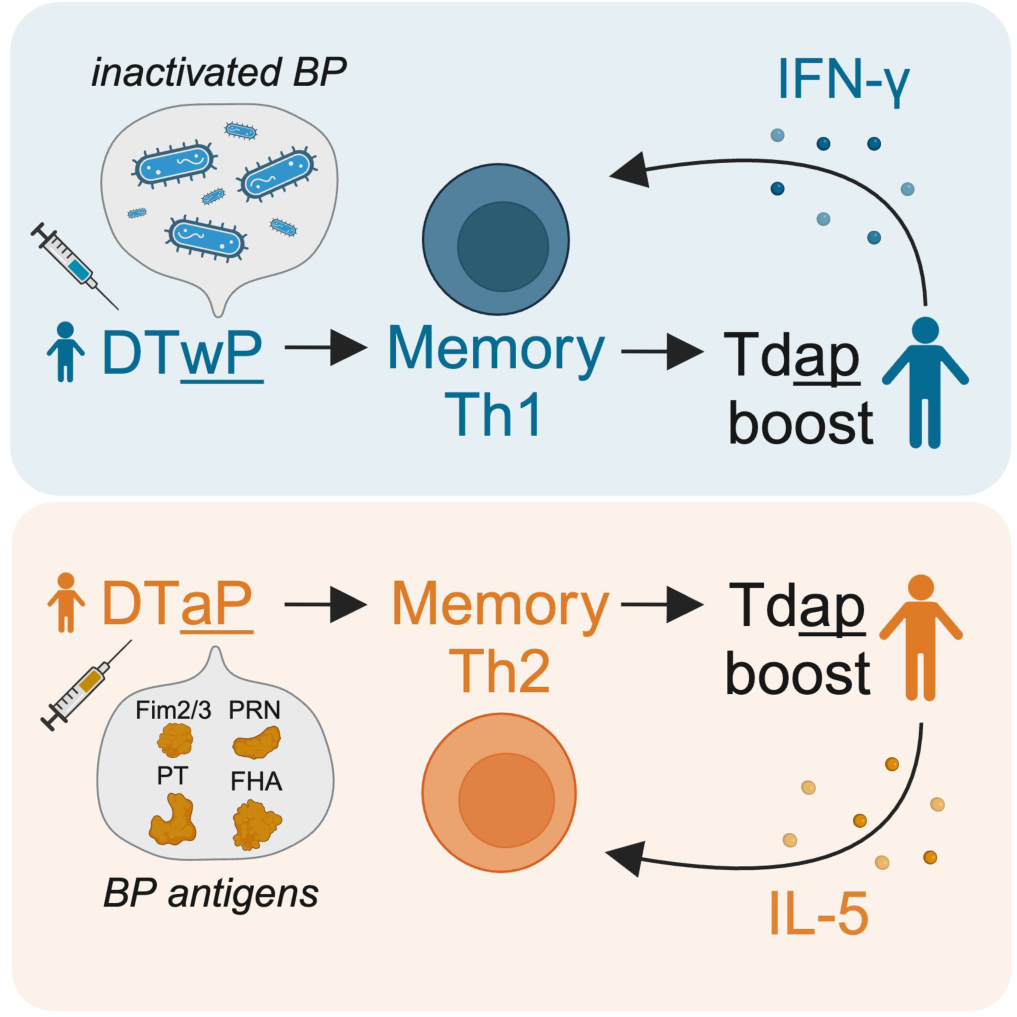
Tdap booster-induced early IFN-γ maintains the Th1 polarized response in DTwP-primed individuals.

A caveat to our finding is that we cannot rule out that type I interferons (IFN-α and IFN-β) rather than IFN-γ are the initial cause of the observed IFN signature on day 1 post-booster. In contrast to IFN-γ, IFN-α and IFN-β can also induce the transcription of genes that have ISRE but lack GAS elements in their promoter. However, since IFN-α, IFN-β, and IFN-γ can all induce transcription of genes containing GAS elements, the possibility cannot be ruled out that both type I and II IFN contribute to the observed booster-induced IFN response that correlates with Th1 polarization. Although a correlation was also found between IFN-γ and Th1 polarization, this IFN-γ could be type I IFN induced as it has been shown that IFN-β stimulates IFN-γ production in human naive CD4^+^ T cells(39). Notably, we did not detect type I interferons at the protein and RNA level in the blood, but that does not rule out their presence in other tissues.

Another limitation of this study is that its design will miss the unmeasured, such as changes in phosphorylation and epigenetics, but also changes in time points and other tissues that were not included. Although our study lacks nasal data, Wilk et al. have described that previous BP infection and wP vaccination, but not aP immunization, lead to the formation of CD69^+^CD4^+^ tissue-resident memory T cells which protect the nose and lungs against BP colonization in C57BL/6 mice(7). This is in line with the findings based on blood describing that the immune response induced by the wP vaccine resembles that induced by natural infection, while the aP-induced immune response is divergent(7, 8). This together suggests that the systemic responses induced by infection and wP vaccination induce overlapping pathways leading to Th1 polarization and, therefore, protection against infection.

Multiple studies report that Th17 cells also play a protective role against BP infection in the nasal mucosa(7, 23). We counted IL-17 secreting cells as a measurement of Th17 polarization using PBMCs and Fluorospot, however, 43 out of the total 98 data points were on the detection limit (data not shown). Therefore, the low sensitivity of the Fluoropot assay to detect IL-17 made these data unsuitable for drawing any meaningful conclusions.

Can Th2 cells be repolarized to Th1 cells to increase the protection against BP infection and duration of immunity? Naive CD4^+^ T cells exhibit significant plasticity in their differentiation into various Th subsets. However, the repolarization capacity of Th1 and Th2 cells seems to disappear once T cells are stimulated multiple times and terminally differentiated(44). This stability is mediated by epigenetic modifications and the expression of lineage-specific transcription factors(45). Additionally, IFN-γ inhibits Th2 polarization and IL-5 inhibits Th1 polarization. Memory Th2 cells retain both antigen specificity and response type, making early vaccination strategy changes more effective in increasing protection against BP infections. However, for individuals already primed with aP vaccines, activating the IFN pathway during booster vaccinations presents a viable alternative to promote Th1 polarization in newly recruited naive T cells, potentially improving the efficacy of current aP vaccines.

How could the IFN pathway be stimulated during vaccination? The first and obvious answer is the use of adjuvants - indispensable vaccine components that stimulate the initiation of an immune response by working as an antigen-delivery system or stimulating Toll-like receptors (TLRs) and other pattern recognition receptors (PRRs)(46, 47). Although the conventional DTaP and DTwP vaccines both contain aluminum salts as adjuvants(48), the exact formulation and type of aluminum salt may vary and be involved in causing the Th1 polarization differences. Aluminum adjuvants are known to induce a Th2 response. FDA-approved adjuvants that induce a Th1 response, such as monophosphoryl lipid A (MPLA) containing adjuvants(49) and the synthetic single-stranded DNA molecule CpG 1018(50) are also available. Changing the pertussis vaccination strategy by switching to or adding an adjuvant that coordinates the immune response to the Th1 phenotype might result in a vaccine that protects longer and better against infection. Interestingly, multiple studies have demonstrated that adding CpG 1018 to Tdap increased protection in mice, with a shift toward Th1, compared to alum-only adjuvanted Tdap(51, 52).

Our studies suggest that changing the current BP vaccination strategy by one that induces IFN-γ production might prevent the lack of robust Th1 responses observed in aP-primed individuals. Further research is needed to verify if targeting IFN-γ during BP vaccination enhances both the longevity of the immune response and protection against infection. Additionally, the specific cell populations responsible for IFN-γ production and their tissue location should be identified. This can first be investigated in the multiple established mouse and non-human primate models, and ultimately in controlled human infection models of BP (53-55).

## Materials and methods

### Study design and human participants

Healthy adults who were primed with either the aP (n=31) or wP (n=30) vaccine during childhood were recruited (**Fig. 1A**). The confirmation of prior receipt of wP or aP containing vaccines for initial priming was accomplished through a multi-faceted approach. Clinical and vaccination records were collected and evaluated for all participants. When vaccination records were unavailable or incomplete, clinical coordinators gathered information about which vaccines, which dates, the number of vaccinations, country of birth, race, and ethnicity through questionnaires. Additionally, birth year was used as a proxy, with participants born in the US before 1995 assumed to have been primed with the wP vaccine in infancy, while those born after 1996 were presumed to have received the aP vaccine. The use of birth year as a proxy for vaccine type is based on the assumption that vaccination practices changed uniformly across the US in 1996, which may not account for regional variations or individual cases that deviated from the norm. All donors were from the San Diego area and were assumed to have followed the recommended vaccination regimen (necessary for enrollment in the California school system). This regimen includes 5 DTaP doses for children under 7 years old, a Tdap booster immunization at 11-12 years followed by a Tdap booster every 10 years. Exclusion criteria included: having received the vaccine within the past 4 years, history of BP infection, current HIV, HBV, or HCV infection, presence of diseases (e.g. significant cardiovascular disease, uncontrolled diabetes, renal disease, liver disease, malignancy, and blood clotting disorder), pregnancy, and in the month before study enrolment: any vaccination, antibiotic or fever (>38°C). Each Tdap booster vaccination (Adacel, Sanofi) dose (0.5 mL) contained tetanus toxoid (TT, 5 Lf), diphtheria toxoid (DT, 2 Lf), the aP antigens: detoxified pertussis toxin (PT, 2.5 μg), and cell surface proteins of BP including filamentous hemagglutinin (FHA, 5 μg), pertactin (PRN, 3 mcg), fimbriae 2/3 (FIM2/3, 5 μg), 1.5 mg aluminum phosphate (0.33 mg aluminum) as the adjuvant, residual formaldehyde (≤5 μg), residual glutaraldehyde (<50 ng) and (0.6% v/v) 2-phenoxyethanol (3.3 mg). Tdap booster vaccination was administered on day 0 and longitudinal blood samples were collected pre-(days -31, -14, 0) and post-booster vaccination (days 1, 3, 7, 14, 28). This study was performed with approvals from the institutional review board at the La Jolla Institute for Immunology (protocol number VD-101). All participants provided written informed consent before donation.

### Overview of performed experiments

Figure 1B shows the experiments that were performed per time point to map the immune response before and after booster vaccination. Characteristics of the study population are shown in **Supplementary Table 1-2** and on the CMI-PB website (https://www.cmi-pb.org/data). Plasma antigen-specific IgG levels including IgG isotypes (IgG1-4), plasma concentrations of 45 different cytokines, and peripheral blood mononuclear cell (PBMC) subset frequencies and transcriptomics experiments were performed previously(40). Here we added measurements of T cell activation and polarization of the same donors at different time points (Fig. 1B**, Supplementary Table 2**).

### Whole blood processing

Whole blood samples (with heparin) were centrifuged at 1850 rpm for 15 min with breaks off. Subsequently, the upper fraction (plasma) was collected and stored at -80°C. Peripheral blood mononuclear cells (PBMC) were isolated by density gradient centrifugation using Ficoll-Paque PLUS (Amersham Biosciences). 35 mL of RPMI 1640 medium (RPMI, Omega Scientific) diluted blood was slowly layered on top of 15 mL Ficoll-Paque PLUS. Sampled were centrifuged at 1850 rpm for 25 min with breaks off. Then, PBMC layers were aspirated and 2 PBMC layers per donor were combined in a new tube together with RPMI. Samples were centrifuged at 1850 rpm for 10 min with a low break. Cell pellets of the same donors were combined and washed with RPMI and centrifuged at 1850 rpm for 10 min with breaks off. Finally, PBMC were counted using trypan blue and a hemocytometer and, after another spin, resuspended in Fetal Bovine Serum (FBS, Gemini) containing 10% DMSO (MilliporeSigma) and stored in a Mr. Frosty cell freezing container overnight at -80°C. The next day, samples were stored in liquid nitrogen until further use.

### AIM assay

Cryopreserved PBMCs were thawed at 37°C for 1 min and added to 10 mL cell culture medium (RPMI 1640 (Corning, 10-041-CM) supplemented with 5% Human Serum AB (GeminiBio, 110-512), 1% Penicillin:Streptomycin solution (GeminiBio, 400-109), and 1% GlutaMAX (Gibco, 35050061)) with 20 uL of Benzonase nuclease (MilliporeSigma). PBMCs were centrifuged at 1400 rpm for 5 min at RT and resuspended in cell culture medium to determine cell concentration and viability using 0.02% Trypan Blue (ThermoFisher) and a hemocytometer. One million PBMCs were seeded in 100uL cell culture medium per well into a 96-well plate (GenClone, 25-221) and were stimulated with 1 of the following 4 stimuli: Tetanus megapool(56) (1 μg/mL), aP megapool(15) (1 μg/mL), 0.3% DMSO (negative control, MilliporeSigma, D2650), and PHA-L (1 μg/mL, positive control, MilliporeSigma, 431784). Stimulated cells were incubated for 18-24 hours at 37°C and 5% CO_2_ and subsequently spun at 1400 rpm for 2 min at RT. Cells were washed twice with PBS (Gibco) and resuspended in 10% FBS in PBS. Then, cells were incubated for 10 min at 4°C. Cells were stained with a 100 uL antibody and dye cocktail (details are shown in **Supplementary Table 5**) for 30 minutes at 4°C in the dark. Stained cells were centrifuged at 1400 rpm for 2 min at 4°C and washed twice with PBS. Cells were resuspended in MACS buffer (2 mM EDTA (Omega) and 0.5% BSA (MilliporeSigma) in PBS at pH 7.0) and transferred to cluster tubes (Corning Life Sciences Plastic, 4401). For compensation, beads (Thermo Fisher, 01-2222-42) were washed twice with PBS and centrifuged at 1400 rpm for 2 min at RT. Beads were distributed in wells and single-stained with 2 uL antibody. Data was acquired using LSR II Flow Cytometer (BD) and analyzed using FlowJo software (v10.10.0). The AIM gating strategy is depicted in **Supplementary Fig. 4**. DMSO values were subtracted from megapool measurements. The median of all DMSO (negative control) measurements (0.012%) was set as the lower detection threshold (LOD) and was added to all measurements before fold change calculations to prevent inflated fold change estimates caused by measurements close to or below the lower detection threshold. In plots where percentages are shown, values below LOD were set to LOD after DMSO subtraction.

### Fluorospot

Two million PBMC were seeded in 500 μL cell culture medium (composition described above). PBMC were incubated with 2 μg/mL aP megapool(15) for 14 days at 37°C and 5% CO_2_. Cells were fed on day 4 with 1 mL of cell culture medium containing 10U/mL IL-2 (Prospec CYT-209B) and 1 mL medium was replaced on days 7 and 11. Fluorospot PVDF plates (Mabtech, 3654-FL-10) were washed with 70% methanol (MilliporeSigma, 154903) and subsequently 3x with sterile dH_2_O. Next, the plates were coated with a capture antibody cocktail containing 10 µg/mL mouse anti-human of each IFN-γ (Mabtech, clone: 1-D1K), IL-5 (Mabtech, clone: TRFK5), and IL-17 (Mabtech, clone: MT44.6) in PBS (Gibco) and incubated overnight at 4°C. The next day, the plates were washed 3x with PBS. Then, plates were blocked by incubating them with cell culture medium for 1 hour at 37°C. After 1 hour, the medium was replaced by a medium containing either 2 μg/mL aP megapool, 0.2% DMSO (negative control, MilliporeSigma, D2650), or 20 μg/mL PHA-L (positive control, MilliporeSigma, 431784). Meanwhile, after 14 days of cell culture, cells were harvested, washed, and counted using a hemocytometer. 2 million cells were added per well in a 1:1 ratio and cultured for 24 hours at 37°C and 5% CO_2_. After 24 hours, the plates were washed 5x with 0.05% Tween20 (MilliporeSigma, PPB005) in PBS. Plates were incubated with the following Mabtech detection antibodies: anti-IFN-γ 7-B6-1-BAM (1:200), anti-IL-5 5A10-WASP (1:200), and 2 μg/mL anti-IL-17 MT504 biotinylated in 0.1% bovine serum albumin (BSA, MilliporeSigma, A3294) in PBS for 2 hours at room temperature (RT). After 2 hours, plates were washed 5x with 0.05% Tween in PBS. Next, plates were incubated with the following Mabtech fluorophore-conjugated detection antibodies: anti-BAM-490, anti-WASP-640, and SA-550 1:200 in 0.1% BSA in PBS for 1 hour at RT, covered from light. After incubation, plates were washed 5x with 0.05% Tween20 in PBS. Then, fluorescence enhancer (Mabtech, 3641-F10) was added and removed after 15 minutes of incubation at RT. Plates were air-dried and measured using a (Mabtech IRIS Fluorospot/ELISpot Reader). A cytokine response was considered positive when the following three criteria were met: 1) eliciting at least 20 spot-forming cells (SFC) per million PBMC after DMSO subtraction, 2) *P*≤0.05 by Student’s t-test or by the Poisson distribution test when comparing the triplicate measurements with the DMSO triplicates, and 3) a greater than 2-fold increase compared to DMSO. If these criteria were met then the SFC per million PBMC value was calculated by averaging the triplicate measurements minus that of DMSO. If these criteria were unfulfilled, the SFC value was set to 20.

### Plasma cytokine concentrations

Plasma samples were randomly distributed on 96 well plates for the absolute quantification of 45 different cytokines (Olink Target 48 Cytokine panel) by Hamilton Health Science (Canada). The Proximity Extension Assay (PEA) technology(57) was used for protein quantification. Briefly, the plasma was incubated with oligonucleotides labeled antibodies targeting the proteins of interest. The oligonucleotides of matched oligonucleotides-antibodies-antigen will bind to each other, enabling amplification and thereby quantification by qPCR. Ct values from the qPCR were used to calculate Normalized Protein eXpression (NPX), a relative quantification unit to report protein expression levels in plasma samples. A standard curve model was established per protein and used to normalize between plates and batches, but also to translate the measured sample NPX values to protein concentrations in pg/mL. More details are described in the Target 48 User Manual (Olink®, v13). Before fold change calculations, the 5% quantile was calculated and added per cytokine to prevent inflated fold change estimates caused by measurements close to or below the lower detection threshold. For cytokines of which the 5% quantile was equal to zero, +1 was applied before fold change calculations.

### Plasma antibody measurements

Pertussis antigen-specific antibody responses were quantified in human plasma by performing an indirect serological assay with xMAP Microspheres (details described in xMAP Cookbook, Luminex 5^th^ edition). Pertussis, Tetanus, and Diphtheria antigens (PT, PRN, Fim2/3, TT, DT (all from List Biological Laboratories), and FHA (MilliporeSigma)) and as a negative control Ovalbumin (OVA, MilliporeSigma) were coupled to uniquely coded beads (xMAP MagPlex Microspheres, Luminex Corporation). PT was inactivated by incubation with 1% formaldehyde (PFA) at 4°C for 10 min. 1% PFA PT and TT were then purified using Zeba spin desalting columns (ThermoFisher). The antigens were coupled with each unique conjugated microsphere at a concentration of 12.5x10^6^ beads/mL using the xMAP Antibody Coupling Kit (Luminex Corporation). Plasma was mixed with a mixture of each conjugated microsphere and the WHO International Standard Human Pertussis antiserum was used as a reference standard (NIBSC, 06/140). Subsequently, the mixtures were washed with 0.05% TWEEN20 in PBS (MilliporeSigma) to exclude non-specific antibodies and targeted antibody responses were detected via anti-human total IgG-PE, IgG1-PE, IgG2-PE, IgG3-PE, and IgG4-PE (all from SouthernBiotech). Antibody details are shown in **Supplementary Table 5**. Samples were subsequently measured on a MAGPIX^®^ instrument (Luminex Corporation) and the log_10_ of the median fluorescent intensity (MFI) values were collected. Measurements were corrected to 0 when the average + 3-fold standard deviation of the background samples (sample dilution buffer) was higher than the measurement.

### RNA sequencing

Per sample, 2 million PBMC were lysed using QIAzol Lysis Reagent (Qiagen). Samples were stored at -80°C until RNA extraction. RNA was extracted using the miRNeasy Mini Kit (Qiagen) including DNase treatment according to the manufacturer’s instructions. The Smart-seq2 protocol was used to perform full-length bulkRNAseq. Briefly, RNA was captured using oligo-poly(dT)-30 primers and reverse transcription was performed using 50-template switching oligos (LNA technologies, Exicon). cDNA was pre-amplified using PCR and purified twice by magnetic beads (volume ratio 0.6x and 0.8x Ampure-XP magnetic beads, Beckman Coulter). cDNA was measured using capillary electrophoresis (Fragment analyzer, Advance analytical) and 0.5 ng of cDNA was used to generate indexed Illumina libraries (Nextera XT library preparation kit, Illumina). Each library’s fragment size was measured by capillary electrophoresis (Fragment analyzer, Advance analytical) and was quantified (Picogreen, Thermofisher). Libraries were pooled at equal molar concentration and sequenced paired-end on an Illumina HiSeq 2500 or NovaSeq 6000 sequencing system to a minimum depth of 15 million reads with each a length of 100 base pairs (S4 flowcell 200 cycle v1.0, Xp workflow; Illumina).

### Bioinformatics RNA sequencing

Raw sequencing reads were aligned to the hg38 genome reference using our in-house pipeline (https://github.com/ndu-UCSD/LJI_RNA_SEQ_PIPELINE_V2). Briefly, FASTQ files were merged and filtered using fastp (v0.20.1), reads were aligned using STAR (v2.7.3a) and further processed using SAMtools (v0.1.19–44428cd), bamCoverage (v3.3.1), and Qualimap (v.2.2.2-dev). From this pipeline, raw counts and transcripts per million (TPM) reads were provided as output. Lowly expressed genes were removed for subsequent analysis (<=10 raw read counts). Differential expression was assessed using the DESeq2 Bioconductor package (v1.36.0)(58) in an R v4.2.1 environment with gene expression called differential with an FDR<0.05. Before fold change calculations, the 5% TPM quantile was calculated per gene and batch year (2021 and 2022) to prevent inflated fold change estimates caused by measurements close to zero. For genes of which the 5% quantile was equal to zero, TPM+0.1 was applied before fold change calculations. Spearman correlation analysis was performed using the correlation package (v0.8.4)(59). Gene set enrichment analysis was performed using Metascape (58) and the Gene Ontology (GO), Reactome, and Kyoto Encyclopedia of Genes and Genomes (KEGG) gene sets(60). Motif enrichment analysis was performed using HOMER findMotifs.pl (v4.11.1) with all genes with at least 10 raw counts as background(61).

## Data availability

Experimental data is publicly available on the CMI-PB website (https://www.cmi-pb.org/data) and PBMC gene expression data are deposited in the Gene Expression Omnibus (GEO) under the accession number GSE274944.

## Acknowledgments

This work was supported by the National Institute of Allergy and Infectious Diseases (NIAID) of the National Institutes of Health (NIH) under award no. U01-AI150753, U01-AI141995 and U19-AI142742. The authors are grateful to all donors who participated in the study and the following La Jolla Institute for Immunology core facilities for their contribution: Clinical Studies, Flow Cytometry, Bioinformatics, and Next Generation Sequencing. Figure 1A-B and Figure 5 were created in BioRender.com.

## Author contributions

Study design: B.P., L.W.; Data collection: F.S.C., L.W., M.A., J.L.; Data analysis and graphical representation: L.W.; Manuscript writing and feedback: L.W., B.P., J.L., R.S.A., S.O., M.K.; Study initiation and supervision: B.P.

## Competing interests

The authors declare no competing interests.

---

**Supplementary Fig. 1.**
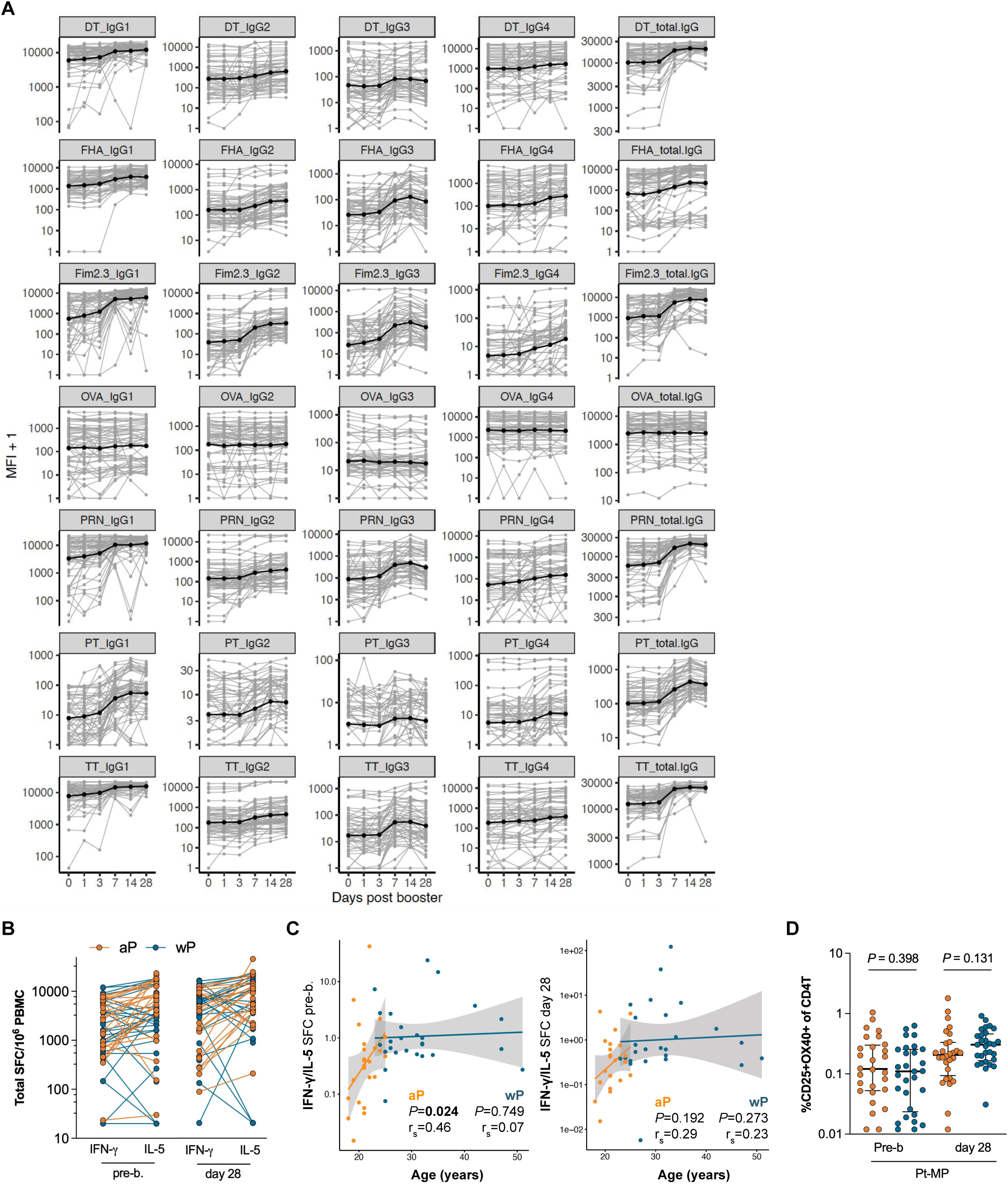
Whole-cell pertussis vaccine priming increases Th1 polarization despite acellular booster vaccination. **A)** Plasma IgG measurements before and after Tdap booster vaccination. Shown are log10-scaled median fluorescence intensities (MFI) of IgG1-4 and total IgG against Tdap antigens (PT, PRN, FHA, FIM2/3, TT, and DT) and the non-Tdap antigen ovalbumin (OVA) for each participant (n=57) and time point (grey). The black lines and points indicate the medians. **B**) IFN-γ and IL-5 producing cells (spot-forming cells, SFC) were measured by Fluorospot after cells were stimulated with aP vaccine antigens for 14 days and derived from blood sampled before and 28 days after Tdap booster vaccination. **C)** Spearman sample correlations between age (years) at booster and T cell polarization data before and 28 days post booster (IFN-γ/IL-5 SFC) per vaccination group. **D)** CD25^+^ and OX40^+^ CD4^+^ T cells were measured before and 28 days after Tdap booster vaccination by flow cytometry and shown as percentage of total CD4^+^ T cells where black lines represent the median with interquartile range. n=28 aP, 29 wP, P-values were calculated by multiple two-tailed Mann-Whitney tests.

**Supplementary Fig. 2.**
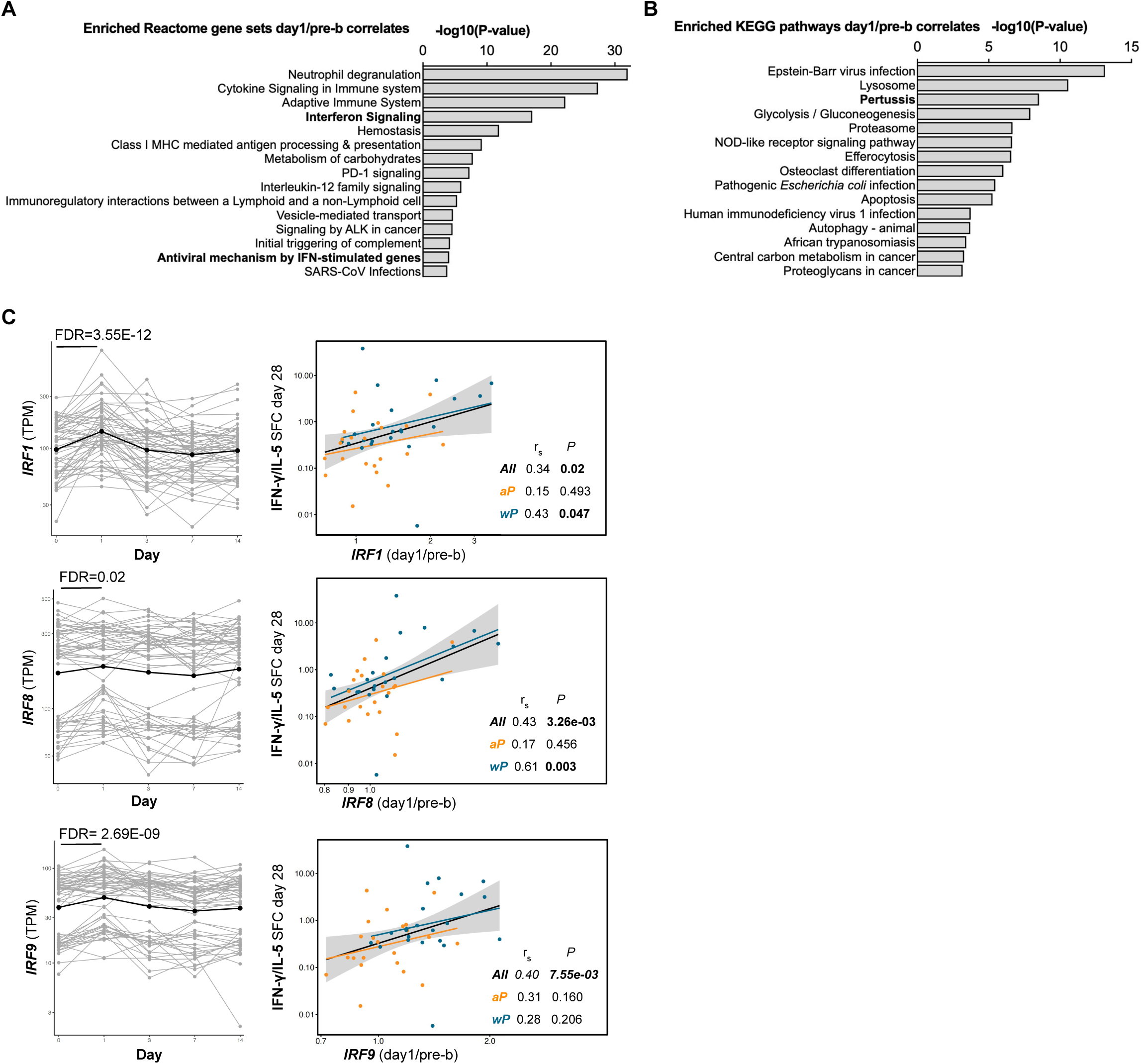
Booster-induced interferon signaling positively correlates with Th1 polarization. **A-B)** The top 15 enriched gene sets identified by **(A)** Reactome or **(B)** Kyoto Encyclopedia of Genes and Genomes (KEGG) gene set enrichment analysis with the 294 DEG (day 1/pre-booster) that also showed a positive correlation with the T cell polarization on day 28. **C)** *IRF1, IRF8* and *IRF9* gene expression (TPM) over time with the black line indicating mean expression and Spearman sample correlation between *IRF’s* (day 1/pre-b) and T cell polarization data 28 days post booster (IFN-γ/IL-5 SFC). Spearman dot plots were created with aP and wP groups combined (“All”) and separately. RNA n=56, correlation analysis n=44. **A-C**) aP and wP-primed individuals were pooled for this analysis.

**Supplementary Fig. 3.**
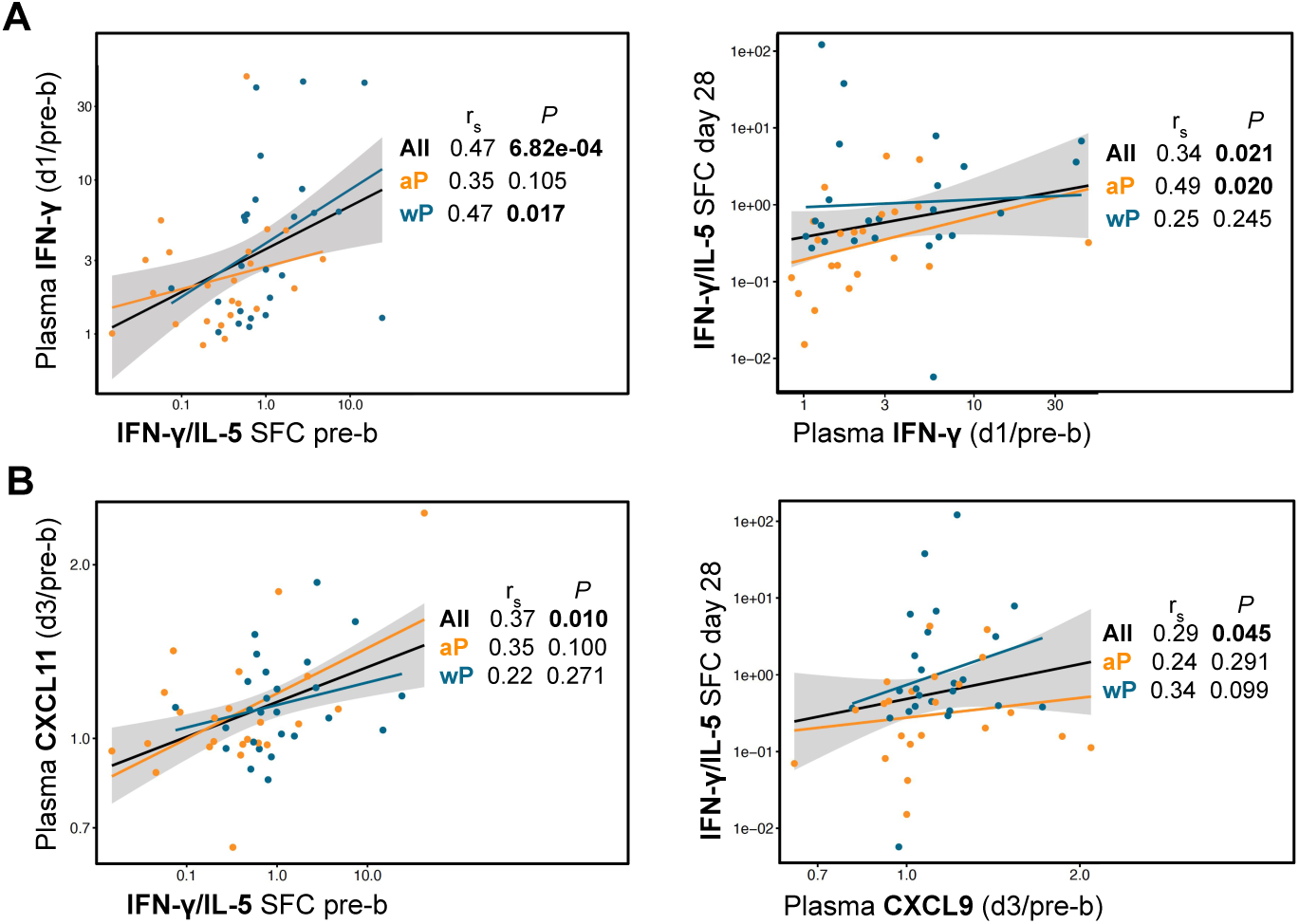
Plasma IFN-γ and IFN-γ initiated chemokines increase post-booster and its changes positively correlate with Th1 polarization. **A-B)** Spearman sample correlation between plasma cytokine changes and T cell polarization pre- and 28 days post booster (IFN-γ/IL-5 SFC). Spearman dot plots were created with aP and wP groups combined (“All”) and separately.

**Supplementary Fig. 4.**
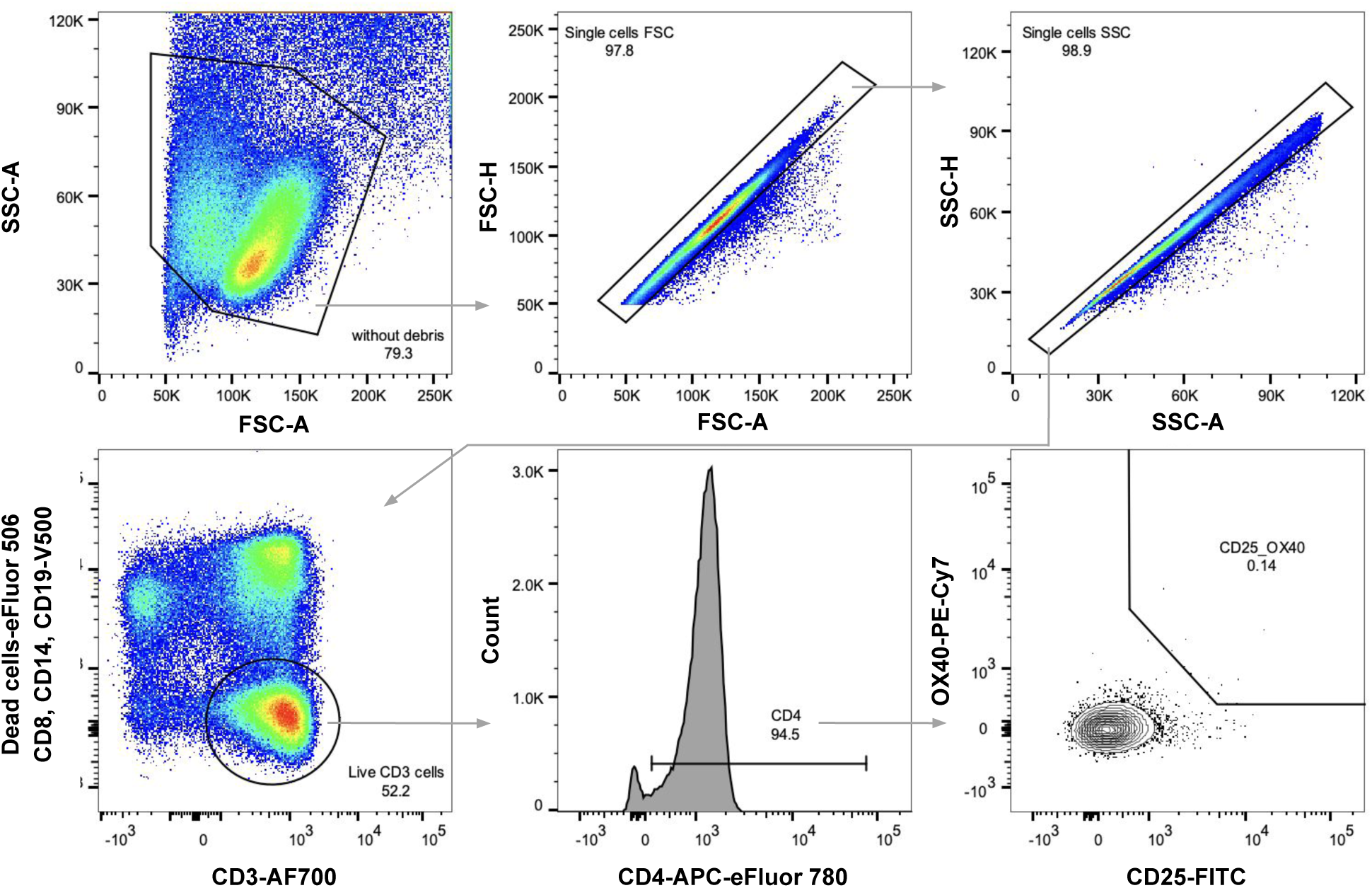
Strategy for flow cytometry gating AIM CD25^+^OX40^+^ CD4^+^ T cells.

## Notes

### Competing Interest Statement

The authors have declared no competing interest.

### Summary of Updates

The manuscript has been improved after receiving helpful reviewer's comments.

