## Supplemental Tables for "Th1 polarization in *Bordetella pertussis* vaccine responses is maintained through a positive feedback loop"

**Supplementary Table 1**. Study population characteristics. Shapiro-Wilk and Mann-Whitney tests were performed to test differences in age at booster between the aP and wP groups. A Chi-square test was used to check if biological sex differed per group.

| | **aP (n=31)** | **wP (n=30)** | ***P*-value** |
|---|---|---|---|
| **Age at booster, mean years ± SD** | 21.1 ± 2.2 | 31.0 ± 7.2 | ***P*<0.001** |
| **Sex, n male (%)** | 10 (32.3%) | 10 (33.3%) | *P*=0.929 |

**Supplementary Table 2**. Performed assays per donor.

| Doras_ID | Subject_ID | Group | AIM | Fluorospot | Plasma antibodies | Plasma cytokines | RNAseq |
|---|---|---|---|---|---|---|---|
| 6797 | 0 | wP | YES | YES | YES | YES | NO |
| 1686 | 2 | wP | YES | YES | YES | YES | NO |
| 2631 | 8 | wP | YES | YES | YES | YES | NO |
| 2726 | 26 | wP | NO | NO | NO | YES | NO |
| 2903 | 45 | aP | NO | NO | NO | YES | NO |
| 3664 | 61 | wP | YES | YES | YES | YES | YES |
| 3802 | 62 | wP | YES | YES | YES | YES | YES |
| 3803 | 63 | wP | YES | YES | YES | YES | YES |
| 3804 | 64 | wP | YES | YES | YES | YES | YES |
| 3806 | 65 | wP | YES | NO | YES | YES | YES |
| 3808 | 66 | wP | YES | YES | YES | YES | YES |
| 3943 | 67 | wP | YES | YES | YES | YES | YES |
| 3944 | 68 | wP | YES | YES | YES | YES | YES |
| 3945 | 69 | wP | YES | YES | YES | YES | YES |
| 3946 | 70 | aP | YES | YES | YES | YES | YES |
| 3947 | 71 | aP | YES | YES | YES | YES | YES |
| 3984 | 72 | wP | YES | YES | YES | YES | YES |
| 3985 | 73 | wP | YES | YES | YES | YES | YES |
| 3986 | 74 | wP | YES | YES | YES | YES | YES |
| 3987 | 75 | aP | YES | YES | YES | YES | YES |
| 3989 | 76 | aP | YES | YES | YES | YES | YES |
| 4016 | 77 | wP | YES | YES | YES | YES | YES |
| 4017 | 78 | wP | YES | YES | YES | YES | YES |
| 4018 | 79 | wP | YES | YES | YES | YES | YES |
| 4019 | 80 | wP | YES | NO | YES | YES | YES |
| 4020 | 81 | wP | YES | YES | YES | YES | YES |
| 4021 | 82 | aP | YES | YES | NO | YES | YES |

| | | | | | | | |
|---|---|---|---|---|---|---|---|
| 4054 | 83 | aP | YES | NO | YES | YES | YES |
| 4055 | 84 | aP | YES | YES | YES | YES | YES |
| 4089 | 85 | aP | YES | NO | YES | YES | YES |
| 4090 | 86 | aP | YES | YES | YES | YES | YES |
| 4091 | 87 | aP | NO | NO | YES | YES | YES |
| 4092 | 88 | aP | NO | NO | NO | YES | YES |
| 4138 | 89 | aP | YES | YES | YES | YES | YES |
| 4139 | 90 | aP | YES | YES | YES | YES | YES |
| 4140 | 91 | aP | YES | YES | YES | YES | YES |
| 4162 | 92 | aP | YES | YES | YES | YES | YES |
| 4163 | 93 | aP | YES | YES | YES | YES | YES |
| 4164 | 94 | aP | YES | YES | YES | YES | YES |
| 4165 | 95 | aP | YES | YES | YES | YES | YES |
| 4166 | 96 | aP | YES | YES | YES | YES | YES |
| 1788 | 97 | wP | YES | YES | YES | YES | YES |
| 4654 | 98 | wP | YES | YES | YES | YES | YES |
| 4656 | 99 | aP | YES | YES | YES | YES | YES |
| 4657 | 100 | aP | YES | NO | YES | YES | YES |
| 6096 | 101 | aP | YES | YES | YES | YES | YES |
| 6097 | 102 | aP | YES | YES | YES | YES | YES |
| 6275 | 103 | wP | YES | YES | YES | YES | YES |
| 6279 | 104 | wP | YES | YES | YES | YES | YES |
| 6280 | 105 | wP | YES | NO | YES | YES | YES |
| 6281 | 106 | aP | YES | YES | YES | YES | YES |
| 6308 | 107 | aP | YES | YES | YES | YES | YES |
| 6385 | 108 | wP | YES | YES | YES | YES | YES |
| 6388 | 109 | wP | YES | YES | YES | YES | YES |
| 6389 | 110 | aP | YES | YES | YES | YES | YES |
| 6482 | 111 | wP | YES | YES | YES | YES | YES |

| | | | | | | | |
|---|---|---|---|---|---|---|---|
| 6483 | 112 | aP | YES | YES | YES | YES | YES |
| 6485 | 114 | wP | YES | YES | YES | YES | YES |
| 6495 | 115 | aP | YES | YES | YES | YES | YES |
| 6679 | 117 | aP | YES | NO | YES | YES | YES |
| 6787 | 118 | aP | YES | YES | YES | YES | YES |

**Supplementary Table 3**. Spearman correlation statistics between gene expression changes (post/pre-b) and T cell polarization (IFN-γ/IL-5 SFC) pre and 28 days post-booster. Significant correlates are shown (P<0.05).

| Parameter 1: Th1 polarization | Parameter 2: Gene (Symbol_ENSEMBL ID) | GEX days | r | *P* | n |
|---|---|---|---|---|---|
| IFN-γ/IL-5 SFC day 28 | GM2A_ENSG00000196743 | day1/pre-b | 0.515 | 0.0003 | 44 |
| IFN-γ/IL-5 SFC day 28 | LAPTM5_ENSG00000162511 | day1/pre-b | 0.512 | 0.0004 | 44 |
| IFN-γ/IL-5 SFC day 28 | TAPBP_ENSG00000231925 | day1/pre-b | 0.508 | 0.0004 | 44 |
| IFN-γ/IL-5 SFC day 28 | MARS1_ENSG00000166986 | day1/pre-b | 0.506 | 0.0005 | 44 |
| IFN-γ/IL-5 SFC pre-b | GBP5_ENSG00000154451 | day1/pre-b | 0.482 | 0.0006 | 47 |
| IFN-γ/IL-5 SFC pre-b | GBP2_ENSG00000162645 | day1/pre-b | 0.478 | 0.0007 | 47 |
| IFN-γ/IL-5 SFC day 28 | HLA.DPB1_ENSG00000223865 | day1/pre-b | 0.488 | 0.0008 | 44 |
| IFN-γ/IL-5 SFC pre-b | STAT1_ENSG00000115415 | day1/pre-b | 0.469 | 0.0009 | 47 |
| IFN-γ/IL-5 SFC day 28 | X_ENSG00000279861 | day1/pre-b | 0.483 | 0.0009 | 44 |
| IFN-γ/IL-5 SFC day 28 | C2_ENSG00000166278 | day1/pre-b | 0.482 | 0.0009 | 44 |
| IFN-γ/IL-5 SFC day 28 | NCSTN_ENSG00000162736 | day1/pre-b | 0.48 | 0.001 | 44 |
| IFN-γ/IL-5 SFC day 28 | P2RX7_ENSG00000089041 | day1/pre-b | 0.479 | 0.001 | 44 |
| IFN-γ/IL-5 SFC day 28 | SUSD6_ENSG00000100647 | day1/pre-b | 0.478 | 0.001 | 44 |
| IFN-γ/IL-5 SFC pre-b | CASP7_ENSG00000165806 | day1/pre-b | 0.462 | 0.0011 | 47 |
| IFN-γ/IL-5 SFC day 28 | ATF5_ENSG00000169136 | day1/pre-b | 0.474 | 0.0012 | 44 |
| IFN-γ/IL-5 SFC day 28 | ADGRE5_ENSG00000123146 | day1/pre-b | 0.47 | 0.0013 | 44 |
| IFN-γ/IL-5 SFC day 28 | APOL4_ENSG00000100336 | day1/pre-b | 0.47 | 0.0013 | 44 |
| IFN-γ/IL-5 SFC day 28 | HLA.DMA_ENSG00000204257 | day1/pre-b | 0.469 | 0.0013 | 44 |
| IFN-γ/IL-5 SFC day 28 | KIF11_ENSG00000138160 | day1/pre-b | 0.467 | 0.0014 | 44 |
| IFN-γ/IL-5 SFC day 28 | USB1_ENSG00000103005 | day1/pre-b | 0.466 | 0.0014 | 44 |
| IFN-γ/IL-5 SFC day 28 | NA_ENSG00000234290 | day1/pre-b | 0.465 | 0.0015 | 44 |
| IFN-γ/IL-5 SFC day 28 | X_ENSG00000254649 | day1/pre-b | 0.464 | 0.0015 | 44 |
| IFN-γ/IL-5 SFC day 28 | TPD52L2_ENSG00000101150 | day1/pre-b | 0.463 | 0.0015 | 44 |
| IFN-γ/IL-5 SFC day 28 | RAB8A_ENSG00000167461 | day1/pre-b | 0.461 | 0.0016 | 44 |
| IFN-γ/IL-5 SFC day 28 | EFTUD2_ENSG00000108883 | day1/pre-b | 0.461 | 0.0017 | 44 |
| IFN-γ/IL-5 SFC pre-b | WARS1_ENSG00000140105 | day1/pre-b | 0.445 | 0.0017 | 47 |
| IFN-γ/IL-5 SFC day 28 | MTCO3P11_ENSG00000237711 | day1/pre-b | 0.459 | 0.0017 | 44 |
| IFN-γ/IL-5 SFC day 28 | MAPRE1_ENSG00000101367 | day1/pre-b | 0.458 | 0.0017 | 44 |
| IFN-γ/IL-5 SFC day 28 | SEC13_ENSG00000157020 | day1/pre-b | 0.458 | 0.0018 | 44 |
| IFN-γ/IL-5 SFC day 28 | MSN_ENSG00000147065 | day1/pre-b | 0.457 | 0.0018 | 44 |
| IFN-γ/IL-5 SFC pre-b | APOL4_ENSG00000100336 | day1/pre-b | 0.441 | 0.0019 | 47 |
| IFN-γ/IL-5 SFC day 28 | GNS_ENSG00000135677 | day1/pre-b | 0.455 | 0.0019 | 44 |
| IFN-γ/IL-5 SFC pre-b | SLAMF8_ENSG00000158714 | day1/pre-b | 0.438 | 0.0021 | 47 |
| IFN-γ/IL-5 SFC day 28 | PRCP_ENSG00000137509 | day1/pre-b | 0.45 | 0.0022 | 44 |
| IFN-γ/IL-5 SFC day 28 | HLA.DMB_ENSG00000242574 | day1/pre-b | 0.45 | 0.0022 | 44 |
| IFN-γ/IL-5 SFC day 28 | MVP_ENSG00000013364 | day1/pre-b | 0.449 | 0.0022 | 44 |

| | | | | | |
|---|---|---|---|---|---|
| IFN-γ/IL-5 SFC day 28 | LIMK2_ENSG00000182541 | day1/pre-b | 0.449 | 0.0022 | 44 |
| IFN-γ/IL-5 SFC pre-b | GBP1_ENSG00000117228 | day1/pre-b | 0.433 | 0.0023 | 47 |
| IFN-γ/IL-5 SFC day 28 | X_ENSG00000258581 | day1/pre-b | 0.446 | 0.0024 | 44 |
| IFN-γ/IL-5 SFC day 28 | RPN1_ENSG00000163902 | day1/pre-b | 0.444 | 0.0025 | 44 |
| IFN-γ/IL-5 SFC pre-b | CYLD.AS1_ENSG00000261644 | day1/pre-b | 0.43 | 0.0025 | 47 |
| IFN-γ/IL-5 SFC day 28 | AKR1A1_ENSG00000117448 | day1/pre-b | 0.443 | 0.0026 | 44 |
| IFN-γ/IL-5 SFC day 28 | SLC35A4_ENSG00000176087 | day1/pre-b | 0.441 | 0.0027 | 44 |
| IFN-γ/IL-5 SFC day 28 | TCN2_ENSG00000185339 | day1/pre-b | 0.441 | 0.0027 | 44 |
| IFN-γ/IL-5 SFC day 28 | STAT2_ENSG00000170581 | day1/pre-b | 0.44 | 0.0028 | 44 |
| IFN-γ/IL-5 SFC day 28 | CASP9_ENSG00000132906 | day1/pre-b | 0.437 | 0.003 | 44 |
| IFN-γ/IL-5 SFC pre-b | PSMB9_ENSG00000240065 | day1/pre-b | 0.423 | 0.003 | 47 |
| IFN-γ/IL-5 SFC day 28 | TPI1_ENSG00000111669 | day1/pre-b | 0.436 | 0.0031 | 44 |
| IFN-γ/IL-5 SFC day 28 | NUCB1_ENSG00000104805 | day1/pre-b | 0.434 | 0.0032 | 44 |
| IFN-γ/IL-5 SFC day 28 | PSAP_ENSG00000197746 | day1/pre-b | 0.434 | 0.0032 | 44 |
| IFN-γ/IL-5 SFC day 28 | IRF8_ENSG00000140968 | day1/pre-b | 0.434 | 0.0033 | 44 |
| IFN-γ/IL-5 SFC day 28 | APOL6_ENSG00000221963 | day1/pre-b | 0.432 | 0.0034 | 44 |
| IFN-γ/IL-5 SFC day 28 | HLA.DQB1_ENSG00000179344 | day1/pre-b | 0.432 | 0.0034 | 44 |
| IFN-γ/IL-5 SFC day 28 | LSP1_ENSG00000130592 | day1/pre-b | 0.43 | 0.0036 | 44 |
| IFN-γ/IL-5 SFC pre-b | TRIM22_ENSG00000132274 | day1/pre-b | 0.417 | 0.0036 | 47 |
| IFN-γ/IL-5 SFC day 28 | ENO1_ENSG00000074800 | day1/pre-b | 0.429 | 0.0037 | 44 |
| IFN-γ/IL-5 SFC day 28 | X_ENSG00000271737 | day1/pre-b | -0.428 | 0.0037 | 44 |
| IFN-γ/IL-5 SFC day 28 | CFL1_ENSG00000172757 | day1/pre-b | 0.427 | 0.0038 | 44 |
| IFN-γ/IL-5 SFC day 28 | RAB20_ENSG00000139832 | day1/pre-b | 0.426 | 0.0039 | 44 |
| IFN-γ/IL-5 SFC pre-b | FAS_ENSG00000026103 | day1/pre-b | 0.411 | 0.0041 | 47 |
| IFN-γ/IL-5 SFC day 28 | CNDP2_ENSG00000133313 | day1/pre-b | 0.423 | 0.0043 | 44 |
| IFN-γ/IL-5 SFC pre-b | APOL1_ENSG00000100342 | day1/pre-b | 0.408 | 0.0044 | 47 |
| IFN-γ/IL-5 SFC day 28 | ADA2_ENSG00000093072 | day1/pre-b | 0.42 | 0.0045 | 44 |
| IFN-γ/IL-5 SFC day 28 | ZBP1_ENSG00000124256 | day1/pre-b | 0.42 | 0.0045 | 44 |
| IFN-γ/IL-5 SFC day 28 | CLK2_ENSG00000176444 | day7/pre-b | -0.419 | 0.0046 | 44 |
| IFN-γ/IL-5 SFC day 28 | TOM1_ENSG00000100284 | day1/pre-b | 0.418 | 0.0047 | 44 |
| IFN-γ/IL-5 SFC day 28 | HDGFL2_ENSG00000167674 | day7/pre-b | -0.418 | 0.0047 | 44 |
| IFN-γ/IL-5 SFC day 28 | C1QA_ENSG00000173372 | day1/pre-b | 0.417 | 0.0049 | 44 |
| IFN-γ/IL-5 SFC day 28 | ARF3_ENSG00000134287 | day1/pre-b | 0.416 | 0.0049 | 44 |
| IFN-γ/IL-5 SFC day 28 | BSG_ENSG00000172270 | day1/pre-b | 0.416 | 0.005 | 44 |
| IFN-γ/IL-5 SFC day 28 | PARVG_ENSG00000138964 | day1/pre-b | 0.413 | 0.0053 | 44 |
| IFN-γ/IL-5 SFC day 28 | DESI1_ENSG00000100418 | day1/pre-b | 0.413 | 0.0053 | 44 |
| IFN-γ/IL-5 SFC day 28 | CD74_ENSG00000019582 | day1/pre-b | 0.413 | 0.0054 | 44 |
| IFN-γ/IL-5 SFC day 28 | IDO1_ENSG00000131203 | day1/pre-b | 0.412 | 0.0055 | 44 |
| IFN-γ/IL-5 SFC pre-b | MYBL1_ENSG00000185697 | day14/pre-b | 0.398 | 0.0056 | 47 |
| IFN-γ/IL-5 SFC day 28 | LILRB1_ENSG00000104972 | day1/pre-b | 0.41 | 0.0057 | 44 |
| IFN-γ/IL-5 SFC pre-b | GBP4_ENSG00000162654 | day1/pre-b | 0.397 | 0.0058 | 47 |

| | | | | | |
|---|---|---|---|---|---|
| IFN-γ/IL-5 SFC day 28 | POLR2E_ENSG00000099817 | day1/pre-b | 0.408 | 0.0059 | 44 |
| IFN-γ/IL-5 SFC day 28 | UBE2L6_ENSG00000156587 | day1/pre-b | 0.407 | 0.0061 | 44 |
| IFN-γ/IL-5 SFC day 28 | PLD3_ENSG00000105223 | day1/pre-b | 0.407 | 0.0061 | 44 |
| IFN-γ/IL-5 SFC day 28 | NAPA_ENSG00000105402 | day1/pre-b | 0.406 | 0.0063 | 44 |
| IFN-γ/IL-5 SFC day 28 | TNFSF13_ENSG00000161955 | day1/pre-b | 0.405 | 0.0063 | 44 |
| IFN-γ/IL-5 SFC day 28 | IFI35_ENSG00000068079 | day1/pre-b | 0.405 | 0.0064 | 44 |
| IFN-γ/IL-5 SFC pre-b | IGHE_ENSG00000211891 | day7/pre-b | -0.391 | 0.0066 | 47 |
| IFN-γ/IL-5 SFC pre-b | FAM30A_ENSG00000226777 | day7/pre-b | -0.391 | 0.0066 | 47 |
| IFN-γ/IL-5 SFC day 28 | HLA.DRB1_ENSG00000196126 | day1/pre-b | 0.402 | 0.0068 | 44 |
| IFN-γ/IL-5 SFC day 28 | PLEK_ENSG00000115956 | day1/pre-b | 0.402 | 0.0068 | 44 |
| IFN-γ/IL-5 SFC day 28 | LRG1_ENSG00000171236 | day1/pre-b | 0.402 | 0.0069 | 44 |
| IFN-γ/IL-5 SFC day 28 | IRF9_ENSG00000213928 | day1/pre-b | 0.401 | 0.007 | 44 |
| IFN-γ/IL-5 SFC day 28 | EIF4E2_ENSG00000135930 | day1/pre-b | 0.4 | 0.0071 | 44 |
| IFN-γ/IL-5 SFC day 28 | SEMA4A_ENSG00000196189 | day1/pre-b | 0.4 | 0.0071 | 44 |
| IFN-γ/IL-5 SFC day 28 | EAF1_ENSG00000144597 | day1/pre-b | 0.4 | 0.0072 | 44 |
| IFN-γ/IL-5 SFC day 28 | FBXO6_ENSG00000116663 | day1/pre-b | 0.4 | 0.0072 | 44 |
| IFN-γ/IL-5 SFC pre-b | IRF9_ENSG00000213928 | day1/pre-b | 0.387 | 0.0073 | 47 |
| IFN-γ/IL-5 SFC day 28 | CD63_ENSG00000135404 | day1/pre-b | 0.397 | 0.0076 | 44 |
| IFN-γ/IL-5 SFC day 28 | DAZAP2_ENSG00000183283 | day1/pre-b | 0.397 | 0.0076 | 44 |
| IFN-γ/IL-5 SFC day 28 | PSMB2_ENSG00000126067 | day1/pre-b | 0.396 | 0.0078 | 44 |
| IFN-γ/IL-5 SFC day 28 | LAP3_ENSG00000002549 | day1/pre-b | 0.395 | 0.0079 | 44 |
| IFN-γ/IL-5 SFC day 28 | SQOR_ENSG00000137767 | day1/pre-b | 0.394 | 0.0081 | 44 |
| IFN-γ/IL-5 SFC day 28 | PTPA_ENSG00000119383 | day1/pre-b | 0.394 | 0.0082 | 44 |
| IFN-γ/IL-5 SFC day 28 | TAPBPL_ENSG00000139192 | day1/pre-b | 0.393 | 0.0083 | 44 |
| IFN-γ/IL-5 SFC day 28 | FANCE_ENSG00000112039 | day14/pre-b | -0.393 | 0.0084 | 44 |
| IFN-γ/IL-5 SFC day 28 | SZRD1_ENSG00000055070 | day1/pre-b | 0.391 | 0.0086 | 44 |
| IFN-γ/IL-5 SFC day 28 | SHKBP1_ENSG00000160410 | day1/pre-b | 0.391 | 0.0087 | 44 |
| IFN-γ/IL-5 SFC day 28 | APOL1_ENSG00000100342 | day1/pre-b | 0.389 | 0.009 | 44 |
| IFN-γ/IL-5 SFC day 28 | GSN_ENSG00000148180 | day1/pre-b | 0.389 | 0.009 | 44 |
| IFN-γ/IL-5 SFC day 28 | TGOLN2_ENSG00000152291 | day1/pre-b | 0.389 | 0.0091 | 44 |
| IFN-γ/IL-5 SFC day 28 | NUP93_ENSG00000102900 | day1/pre-b | 0.389 | 0.0091 | 44 |
| IFN-γ/IL-5 SFC day 28 | SHTN1_ENSG00000187164 | day1/pre-b | 0.388 | 0.0092 | 44 |
| IFN-γ/IL-5 SFC day 28 | GRN_ENSG00000030582 | day1/pre-b | 0.388 | 0.0093 | 44 |
| IFN-γ/IL-5 SFC day 28 | CTSH_ENSG00000103811 | day1/pre-b | 0.388 | 0.0093 | 44 |
| IFN-γ/IL-5 SFC pre-b | SERPING1_ENSG00000149131 | day1/pre-b | 0.375 | 0.0093 | 47 |
| IFN-γ/IL-5 SFC day 28 | ACLY_ENSG00000131473 | day1/pre-b | 0.388 | 0.0093 | 44 |
| IFN-γ/IL-5 SFC day 28 | LGALS9_ENSG00000168961 | day1/pre-b | 0.388 | 0.0093 | 44 |
| IFN-γ/IL-5 SFC day 28 | GABARAP_ENSG00000170296 | day1/pre-b | 0.387 | 0.0094 | 44 |
| IFN-γ/IL-5 SFC day 28 | PSMD2_ENSG00000175166 | day1/pre-b | 0.387 | 0.0094 | 44 |
| IFN-γ/IL-5 SFC day 28 | UBE2D3_ENSG00000109332 | day1/pre-b | 0.386 | 0.0096 | 44 |
| IFN-γ/IL-5 SFC day 28 | RAB7A_ENSG00000075785 | day1/pre-b | 0.386 | 0.0097 | 44 |

| | | | | | |
|---|---|---|---|---|---|
| IFN-γ/IL-5 SFC day 28 | NAGA_ENSG00000198951 | day1/pre-b | 0.385 | 0.0099 | 44 |
| IFN-γ/IL-5 SFC day 28 | DBNL_ENSG00000136279 | day1/pre-b | 0.384 | 0.01 | 44 |
| IFN-γ/IL-5 SFC day 28 | LILRB4_ENSG00000186818 | day1/pre-b | 0.384 | 0.0101 | 44 |
| IFN-γ/IL-5 SFC pre-b | IGKV2OR22.4_ENSG00000253691 | day7/pre-b | -0.371 | 0.0103 | 47 |
| IFN-γ/IL-5 SFC day 28 | JARID2_ENSG00000008083 | day1/pre-b | 0.382 | 0.0105 | 44 |
| IFN-γ/IL-5 SFC day 28 | RNF26_ENSG00000173456 | day1/pre-b | 0.381 | 0.0106 | 44 |
| IFN-γ/IL-5 SFC day 28 | DTX3L_ENSG00000163840 | day1/pre-b | 0.381 | 0.0108 | 44 |
| IFN-γ/IL-5 SFC day 28 | FXR2_ENSG00000129245 | day7/pre-b | -0.381 | 0.0108 | 44 |
| IFN-γ/IL-5 SFC day 28 | MELK_ENSG00000165304 | day7/pre-b | 0.381 | 0.0108 | 44 |
| IFN-γ/IL-5 SFC pre-b | HLA.DRB5_ENSG00000198502 | day1/pre-b | 0.368 | 0.0108 | 47 |
| IFN-γ/IL-5 SFC day 28 | FAM53C_ENSG00000120709 | day1/pre-b | 0.38 | 0.0109 | 44 |
| IFN-γ/IL-5 SFC day 28 | EIF4H_ENSG00000106682 | day1/pre-b | 0.38 | 0.011 | 44 |
| IFN-γ/IL-5 SFC day 28 | PEPD_ENSG00000124299 | day1/pre-b | 0.38 | 0.011 | 44 |
| IFN-γ/IL-5 SFC day 28 | MYOF_ENSG00000138119 | day1/pre-b | 0.379 | 0.0111 | 44 |
| IFN-γ/IL-5 SFC day 28 | CUL1_ENSG00000055130 | day1/pre-b | 0.379 | 0.0113 | 44 |
| IFN-γ/IL-5 SFC day 28 | PTPN6_ENSG00000111679 | day1/pre-b | 0.378 | 0.0115 | 44 |
| IFN-γ/IL-5 SFC day 28 | IGHV2.5_ENSG00000211937 | day7/pre-b | 0.377 | 0.0115 | 44 |
| IFN-γ/IL-5 SFC day 28 | BISPR_ENSG00000282851 | day1/pre-b | 0.377 | 0.0116 | 44 |
| IFN-γ/IL-5 SFC pre-b | PSME2_ENSG00000100911 | day1/pre-b | 0.364 | 0.0118 | 47 |
| IFN-γ/IL-5 SFC day 28 | TFE3_ENSG00000068323 | day1/pre-b | 0.376 | 0.0118 | 44 |
| IFN-γ/IL-5 SFC day 28 | SLC31A1_ENSG00000136868 | day1/pre-b | 0.376 | 0.0118 | 44 |
| IFN-γ/IL-5 SFC day 28 | IGHE_ENSG00000211891 | day7/pre-b | -0.376 | 0.0119 | 44 |
| IFN-γ/IL-5 SFC day 28 | WARS1_ENSG00000140105 | day1/pre-b | 0.376 | 0.012 | 44 |
| IFN-γ/IL-5 SFC day 28 | MPEG1_ENSG00000197629 | day1/pre-b | 0.376 | 0.012 | 44 |
| IFN-γ/IL-5 SFC day 28 | PGAM1_ENSG00000171314 | day1/pre-b | 0.376 | 0.012 | 44 |
| IFN-γ/IL-5 SFC pre-b | UBE2L6_ENSG00000156587 | day1/pre-b | 0.363 | 0.012 | 47 |
| IFN-γ/IL-5 SFC day 28 | MAPK1IP1L_ENSG00000168175 | day1/pre-b | 0.375 | 0.0121 | 44 |
| IFN-γ/IL-5 SFC day 28 | MYD88_ENSG00000172936 | day1/pre-b | 0.375 | 0.0122 | 44 |
| IFN-γ/IL-5 SFC day 28 | AAR2_ENSG00000131043 | day1/pre-b | 0.375 | 0.0122 | 44 |
| IFN-γ/IL-5 SFC day 28 | UPP1_ENSG00000183696 | day1/pre-b | 0.375 | 0.0122 | 44 |
| IFN-γ/IL-5 SFC day 28 | SERPING1_ENSG00000149131 | day1/pre-b | 0.374 | 0.0123 | 44 |
| IFN-γ/IL-5 SFC day 28 | NECAP1_ENSG00000089818 | day1/pre-b | 0.374 | 0.0124 | 44 |
| IFN-γ/IL-5 SFC pre-b | FBXO6_ENSG00000116663 | day1/pre-b | 0.361 | 0.0126 | 47 |
| IFN-γ/IL-5 SFC pre-b | IGLV2.8_ENSG00000278196 | day7/pre-b | -0.361 | 0.0127 | 47 |
| IFN-γ/IL-5 SFC day 28 | ANKRD22_ENSG00000152766 | day1/pre-b | 0.372 | 0.013 | 44 |
| IFN-γ/IL-5 SFC day 28 | NLRC5_ENSG00000140853 | day1/pre-b | 0.371 | 0.0132 | 44 |
| IFN-γ/IL-5 SFC day 28 | SLC6A12_ENSG00000111181 | day1/pre-b | 0.371 | 0.0133 | 44 |
| IFN-γ/IL-5 SFC pre-b | IRF1_ENSG00000125347 | day1/pre-b | 0.358 | 0.0134 | 47 |
| IFN-γ/IL-5 SFC day 28 | TRIM26_ENSG00000234127 | day1/pre-b | 0.37 | 0.0135 | 44 |
| IFN-γ/IL-5 SFC day 28 | TNFRSF10B_ENSG00000120889 | day1/pre-b | 0.37 | 0.0135 | 44 |
| IFN-γ/IL-5 SFC day 28 | AOAH_ENSG00000136250 | day1/pre-b | 0.369 | 0.0137 | 44 |

| | | | | | |
|---|---|---|---|---|---|
| IFN-γ/IL-5 SFC day 28 | X_ENSG00000261025 | day1/pre-b | -0.367 | 0.0143 | 44 |
| IFN-γ/IL-5 SFC day 28 | WBP11_ENSG00000084463 | day1/pre-b | 0.367 | 0.0143 | 44 |
| IFN-γ/IL-5 SFC day 28 | RNH1_ENSG00000023191 | day1/pre-b | 0.367 | 0.0144 | 44 |
| IFN-γ/IL-5 SFC pre-b | HLA.DPA1_ENSG00000231389 | day1/pre-b | 0.354 | 0.0146 | 47 |
| IFN-γ/IL-5 SFC day 28 | NELFB_ENSG00000188986 | day7/pre-b | -0.366 | 0.0146 | 44 |
| IFN-γ/IL-5 SFC day 28 | KXD1_ENSG00000105700 | day1/pre-b | 0.365 | 0.0147 | 44 |
| IFN-γ/IL-5 SFC day 28 | LCP1_ENSG00000136167 | day1/pre-b | 0.365 | 0.0148 | 44 |
| IFN-γ/IL-5 SFC day 28 | KLF10_ENSG00000155090 | day1/pre-b | 0.365 | 0.0148 | 44 |
| IFN-γ/IL-5 SFC pre-b | GSDMD_ENSG00000104518 | day1/pre-b | 0.353 | 0.0148 | 47 |
| IFN-γ/IL-5 SFC pre-b | HLA.DRA_ENSG00000204287 | day1/pre-b | 0.353 | 0.0151 | 47 |
| IFN-γ/IL-5 SFC day 28 | GYG1_ENSG00000163754 | day1/pre-b | 0.364 | 0.0152 | 44 |
| IFN-γ/IL-5 SFC pre-b | GAB1_ENSG00000109458 | day7/pre-b | 0.351 | 0.0154 | 47 |
| IFN-γ/IL-5 SFC pre-b | ANKRD22_ENSG00000152766 | day1/pre-b | 0.351 | 0.0155 | 47 |
| IFN-γ/IL-5 SFC day 28 | FCGRT_ENSG00000104870 | day1/pre-b | 0.363 | 0.0156 | 44 |
| IFN-γ/IL-5 SFC day 28 | LINC00570_ENSG00000224177 | day7/pre-b | -0.361 | 0.0159 | 44 |
| IFN-γ/IL-5 SFC pre-b | RIOK3_ENSG00000101782 | day7/pre-b | 0.35 | 0.016 | 47 |
| IFN-γ/IL-5 SFC day 28 | RAB5C_ENSG00000108774 | day1/pre-b | 0.361 | 0.0161 | 44 |
| IFN-γ/IL-5 SFC day 28 | PPCDC_ENSG00000138621 | day1/pre-b | 0.361 | 0.0162 | 44 |
| IFN-γ/IL-5 SFC pre-b | ETV7_ENSG00000010030 | day1/pre-b | 0.349 | 0.0162 | 47 |
| IFN-γ/IL-5 SFC day 28 | IL1RN_ENSG00000136689 | day1/pre-b | 0.36 | 0.0163 | 44 |
| IFN-γ/IL-5 SFC day 28 | SLC3A2_ENSG00000168003 | day1/pre-b | 0.36 | 0.0164 | 44 |
| IFN-γ/IL-5 SFC day 28 | STXBP2_ENSG00000076944 | day1/pre-b | 0.36 | 0.0164 | 44 |
| IFN-γ/IL-5 SFC day 28 | PSMB6_ENSG00000142507 | day1/pre-b | 0.36 | 0.0165 | 44 |
| IFN-γ/IL-5 SFC day 28 | VPS18_ENSG00000104142 | day1/pre-b | 0.359 | 0.0166 | 44 |
| IFN-γ/IL-5 SFC day 28 | SHISA5_ENSG00000164054 | day1/pre-b | 0.359 | 0.0167 | 44 |
| IFN-γ/IL-5 SFC day 28 | JMJD6_ENSG00000070495 | day1/pre-b | 0.359 | 0.0167 | 44 |
| IFN-γ/IL-5 SFC day 28 | NA_ENSG00000112096 | day1/pre-b | 0.359 | 0.0167 | 44 |
| IFN-γ/IL-5 SFC day 28 | SDHA_ENSG00000073578 | day1/pre-b | 0.358 | 0.0169 | 44 |
| IFN-γ/IL-5 SFC pre-b | GLS_ENSG00000115419 | day14/pre-b | 0.347 | 0.017 | 47 |
| IFN-γ/IL-5 SFC day 28 | FCN1_ENSG00000085265 | day1/pre-b | 0.358 | 0.017 | 44 |
| IFN-γ/IL-5 SFC day 28 | PGK1_ENSG00000102144 | day1/pre-b | 0.358 | 0.017 | 44 |
| IFN-γ/IL-5 SFC pre-b | BDNF_ENSG00000176697 | day7/pre-b | 0.346 | 0.0172 | 47 |
| IFN-γ/IL-5 SFC day 28 | CTSB_ENSG00000164733 | day1/pre-b | 0.357 | 0.0173 | 44 |
| IFN-γ/IL-5 SFC day 28 | CAP1_ENSG00000131236 | day1/pre-b | 0.357 | 0.0173 | 44 |
| IFN-γ/IL-5 SFC pre-b | PSMB10_ENSG00000205220 | day1/pre-b | 0.346 | 0.0174 | 47 |
| IFN-γ/IL-5 SFC day 28 | RPL32P11_ENSG00000213872 | day1/pre-b | -0.356 | 0.0177 | 44 |
| IFN-γ/IL-5 SFC day 28 | DRAP1_ENSG00000175550 | day1/pre-b | 0.356 | 0.0177 | 44 |
| IFN-γ/IL-5 SFC day 28 | HSD3B7_ENSG00000099377 | day1/pre-b | 0.356 | 0.0178 | 44 |
| IFN-γ/IL-5 SFC day 28 | JDP2_ENSG00000140044 | day1/pre-b | 0.355 | 0.0179 | 44 |
| IFN-γ/IL-5 SFC pre-b | LAP3_ENSG00000002549 | day1/pre-b | 0.344 | 0.0179 | 47 |
| IFN-γ/IL-5 SFC pre-b | REC8_ENSG00000100918 | day1/pre-b | 0.344 | 0.018 | 47 |

| | | | | | |
|---|---|---|---|---|---|
| IFN-γ/IL-5 SFC day 28 | ZNF181_ENSG00000197841 | day1/pre-b | -0.355 | 0.018 | 44 |
| IFN-γ/IL-5 SFC pre-b | EPSTI1_ENSG00000133106 | day1/pre-b | 0.343 | 0.0184 | 47 |
| IFN-γ/IL-5 SFC pre-b | HLA.DRB1_ENSG00000196126 | day1/pre-b | 0.343 | 0.0184 | 47 |
| IFN-γ/IL-5 SFC day 28 | TRAFD1_ENSG00000135148 | day1/pre-b | 0.354 | 0.0184 | 44 |
| IFN-γ/IL-5 SFC day 28 | BAK1_ENSG00000030110 | day1/pre-b | 0.354 | 0.0184 | 44 |
| IFN-γ/IL-5 SFC pre-b | BATF2_ENSG00000168062 | day1/pre-b | 0.342 | 0.0185 | 47 |
| IFN-γ/IL-5 SFC pre-b | PLK1_ENSG00000166851 | day7/pre-b | -0.342 | 0.0186 | 47 |
| IFN-γ/IL-5 SFC day 28 | ITGB2_ENSG00000160255 | day1/pre-b | 0.353 | 0.0186 | 44 |
| IFN-γ/IL-5 SFC day 28 | LCP2_ENSG00000043462 | day1/pre-b | 0.353 | 0.0186 | 44 |
| IFN-γ/IL-5 SFC day 28 | DCP1A_ENSG00000272886 | day1/pre-b | 0.353 | 0.0186 | 44 |
| IFN-γ/IL-5 SFC day 28 | SEC14L1_ENSG00000129657 | day1/pre-b | 0.352 | 0.019 | 44 |
| IFN-γ/IL-5 SFC day 28 | OGDH_ENSG00000105953 | day1/pre-b | 0.352 | 0.0191 | 44 |
| IFN-γ/IL-5 SFC day 28 | X_ENSG00000249138 | day1/pre-b | 0.352 | 0.0191 | 44 |
| IFN-γ/IL-5 SFC day 28 | NQO2_ENSG00000124588 | day1/pre-b | 0.352 | 0.0192 | 44 |
| IFN-γ/IL-5 SFC pre-b | PSMB3_ENSG00000277791 | day1/pre-b | 0.34 | 0.0193 | 47 |
| IFN-γ/IL-5 SFC day 28 | SECTM1_ENSG00000141574 | day1/pre-b | 0.352 | 0.0193 | 44 |
| IFN-γ/IL-5 SFC day 28 | FGR_ENSG00000000938 | day1/pre-b | 0.352 | 0.0193 | 44 |
| IFN-γ/IL-5 SFC day 28 | APOBEC3A_ENSG00000128383 | day1/pre-b | 0.351 | 0.0194 | 44 |
| IFN-γ/IL-5 SFC pre-b | DNAI4_ENSG00000152763 | day1/pre-b | 0.339 | 0.0199 | 47 |
| IFN-γ/IL-5 SFC day 28 | LDHA_ENSG00000134333 | day1/pre-b | 0.35 | 0.02 | 44 |
| IFN-γ/IL-5 SFC day 28 | FARP2_ENSG00000006607 | day1/pre-b | 0.35 | 0.02 | 44 |
| IFN-γ/IL-5 SFC pre-b | ORAI3_ENSG00000175938 | day7/pre-b | -0.338 | 0.02 | 47 |
| IFN-γ/IL-5 SFC pre-b | ZNF567.DT_ENSG00000225975 | day1/pre-b | 0.338 | 0.0201 | 47 |
| IFN-γ/IL-5 SFC day 28 | LTA4H_ENSG00000111144 | day1/pre-b | 0.349 | 0.0201 | 44 |
| IFN-γ/IL-5 SFC day 28 | GK_ENSG00000198814 | day1/pre-b | 0.349 | 0.0204 | 44 |
| IFN-γ/IL-5 SFC day 28 | LILRB3_ENSG00000204577 | day1/pre-b | 0.348 | 0.0205 | 44 |
| IFN-γ/IL-5 SFC day 28 | FCGR1A_ENSG00000150337 | day1/pre-b | 0.348 | 0.0207 | 44 |
| IFN-γ/IL-5 SFC day 28 | GLUL_ENSG00000135821 | day1/pre-b | 0.347 | 0.021 | 44 |
| IFN-γ/IL-5 SFC day 28 | DEXI_ENSG00000182108 | day7/pre-b | 0.347 | 0.021 | 44 |
| IFN-γ/IL-5 SFC pre-b | GBP1P1_ENSG00000225492 | day1/pre-b | 0.336 | 0.021 | 47 |
| IFN-γ/IL-5 SFC day 28 | PILRA_ENSG00000085514 | day1/pre-b | 0.346 | 0.0213 | 44 |
| IFN-γ/IL-5 SFC day 28 | TNFSF13B_ENSG00000102524 | day1/pre-b | 0.346 | 0.0214 | 44 |
| IFN-γ/IL-5 SFC day 28 | GBP4_ENSG00000162654 | day1/pre-b | 0.346 | 0.0215 | 44 |
| IFN-γ/IL-5 SFC pre-b | PSME1_ENSG00000092010 | day1/pre-b | 0.334 | 0.0217 | 47 |
| IFN-γ/IL-5 SFC day 28 | IGLV3.9_ENSG00000211670 | day7/pre-b | 0.345 | 0.0217 | 44 |
| IFN-γ/IL-5 SFC day 28 | HLA.DRA_ENSG00000204287 | day1/pre-b | 0.345 | 0.0219 | 44 |
| IFN-γ/IL-5 SFC day 28 | CFP_ENSG00000126759 | day1/pre-b | 0.345 | 0.0219 | 44 |
| IFN-γ/IL-5 SFC day 28 | PKM_ENSG00000067225 | day1/pre-b | 0.345 | 0.022 | 44 |
| IFN-γ/IL-5 SFC day 28 | PPP4C_ENSG00000149923 | day1/pre-b | 0.344 | 0.0222 | 44 |
| IFN-γ/IL-5 SFC pre-b | CUL1_ENSG00000055130 | day1/pre-b | 0.333 | 0.0222 | 47 |
| IFN-γ/IL-5 SFC day 28 | HSPA5_ENSG00000044574 | day1/pre-b | 0.343 | 0.0225 | 44 |

| | | | | | |
|---|---|---|---|---|---|
| IFN-γ/IL-5 SFC day 28 | TRAPPC14_ENSG00000146826 | day7/pre-b | -0.343 | 0.0226 | 44 |
| IFN-γ/IL-5 SFC pre-b | PADI6_ENSG00000276747 | day1/pre-b | 0.332 | 0.0227 | 47 |
| IFN-γ/IL-5 SFC day 28 | ARF1_ENSG00000143761 | day1/pre-b | 0.343 | 0.0227 | 44 |
| IFN-γ/IL-5 SFC day 28 | SIRPG.AS1_ENSG00000237914 | day1/pre-b | 0.343 | 0.0227 | 44 |
| IFN-γ/IL-5 SFC day 28 | PSMB3_ENSG00000277791 | day1/pre-b | 0.343 | 0.0228 | 44 |
| IFN-γ/IL-5 SFC pre-b | HLA.DQB1_ENSG00000179344 | day1/pre-b | 0.331 | 0.023 | 47 |
| IFN-γ/IL-5 SFC day 28 | MCTP1_ENSG00000175471 | day1/pre-b | 0.342 | 0.023 | 44 |
| IFN-γ/IL-5 SFC day 28 | EIF3B_ENSG00000106263 | day7/pre-b | -0.342 | 0.0231 | 44 |
| IFN-γ/IL-5 SFC day 28 | CIBAR1_ENSG00000188343 | day1/pre-b | 0.342 | 0.0232 | 44 |
| IFN-γ/IL-5 SFC pre-b | HBA2_ENSG00000188536 | day7/pre-b | 0.331 | 0.0232 | 47 |
| IFN-γ/IL-5 SFC day 28 | AP2M1_ENSG00000161203 | day1/pre-b | 0.341 | 0.0233 | 44 |
| IFN-γ/IL-5 SFC day 28 | OTUB1_ENSG00000167770 | day1/pre-b | 0.341 | 0.0236 | 44 |
| IFN-γ/IL-5 SFC pre-b | VAMP5_ENSG00000168899 | day1/pre-b | 0.329 | 0.0238 | 47 |
| IFN-γ/IL-5 SFC pre-b | GK4P_ENSG00000178146 | day1/pre-b | 0.329 | 0.0239 | 47 |
| IFN-γ/IL-5 SFC day 28 | PPP1CA_ENSG00000172531 | day1/pre-b | 0.34 | 0.024 | 44 |
| IFN-γ/IL-5 SFC day 28 | RBCK1_ENSG00000125826 | day1/pre-b | 0.34 | 0.0241 | 44 |
| IFN-γ/IL-5 SFC day 28 | SLC7A7_ENSG00000155465 | day1/pre-b | 0.338 | 0.0248 | 44 |
| IFN-γ/IL-5 SFC day 28 | IRF1_ENSG00000125347 | day1/pre-b | 0.338 | 0.0248 | 44 |
| IFN-γ/IL-5 SFC day 28 | X_ENSG00000216775 | day7/pre-b | 0.338 | 0.025 | 44 |
| IFN-γ/IL-5 SFC day 28 | CAPG_ENSG00000042493 | day1/pre-b | 0.337 | 0.0252 | 44 |
| IFN-γ/IL-5 SFC pre-b | FANCE_ENSG00000112039 | day14/pre-b | -0.326 | 0.0252 | 47 |
| IFN-γ/IL-5 SFC day 28 | CTNNAL1_ENSG00000119326 | day7/pre-b | -0.337 | 0.0254 | 44 |
| IFN-γ/IL-5 SFC day 28 | LAMP2_ENSG00000005893 | day1/pre-b | 0.337 | 0.0255 | 44 |
| IFN-γ/IL-5 SFC day 28 | ANXA5_ENSG00000164111 | day1/pre-b | 0.336 | 0.0256 | 44 |
| IFN-γ/IL-5 SFC day 28 | CYLD.AS1_ENSG00000261644 | day1/pre-b | 0.336 | 0.0256 | 44 |
| IFN-γ/IL-5 SFC day 28 | TMEM179B_ENSG00000185475 | day1/pre-b | 0.336 | 0.0257 | 44 |
| IFN-γ/IL-5 SFC pre-b | MTHFD2_ENSG00000065911 | day1/pre-b | 0.325 | 0.0258 | 47 |
| IFN-γ/IL-5 SFC day 28 | SAT2_ENSG00000141504 | day1/pre-b | 0.336 | 0.0258 | 44 |
| IFN-γ/IL-5 SFC day 28 | ZNF394_ENSG00000160908 | day1/pre-b | 0.336 | 0.0258 | 44 |
| IFN-γ/IL-5 SFC day 28 | CYBB_ENSG00000165168 | day1/pre-b | 0.336 | 0.0258 | 44 |
| IFN-γ/IL-5 SFC day 28 | CES1_ENSG00000198848 | day1/pre-b | 0.336 | 0.0259 | 44 |
| IFN-γ/IL-5 SFC pre-b | HCAR2_ENSG00000182782 | day1/pre-b | 0.325 | 0.026 | 47 |
| IFN-γ/IL-5 SFC day 28 | ICAM1_ENSG00000090339 | day1/pre-b | 0.335 | 0.0262 | 44 |
| IFN-γ/IL-5 SFC day 28 | PSME2_ENSG00000100911 | day1/pre-b | 0.335 | 0.0264 | 44 |
| IFN-γ/IL-5 SFC day 28 | LILRA2_ENSG00000239998 | day1/pre-b | 0.334 | 0.0266 | 44 |
| IFN-γ/IL-5 SFC day 28 | HCK_ENSG00000101336 | day1/pre-b | 0.334 | 0.0269 | 44 |
| IFN-γ/IL-5 SFC day 28 | NMI_ENSG00000123609 | day1/pre-b | 0.333 | 0.027 | 44 |
| IFN-γ/IL-5 SFC day 28 | MX2_ENSG00000183486 | day1/pre-b | 0.333 | 0.0271 | 44 |
| IFN-γ/IL-5 SFC day 28 | MPZL2_ENSG00000149573 | day1/pre-b | 0.333 | 0.0272 | 44 |
| IFN-γ/IL-5 SFC pre-b | ZNF383_ENSG00000188283 | day1/pre-b | 0.322 | 0.0272 | 47 |
| IFN-γ/IL-5 SFC day 28 | CORO1B_ENSG00000172725 | day1/pre-b | 0.332 | 0.0275 | 44 |

| | | | | | |
|---|---|---|---|---|---|
| IFN-γ/IL-5 SFC day 28 | ADAM17_ENSG00000151694 | day1/pre-b | 0.332 | 0.0279 | 44 |
| IFN-γ/IL-5 SFC day 28 | CASP7_ENSG00000165806 | day1/pre-b | 0.332 | 0.0279 | 44 |
| IFN-γ/IL-5 SFC day 28 | STAT3_ENSG00000168610 | day1/pre-b | 0.331 | 0.028 | 44 |
| IFN-γ/IL-5 SFC day 28 | EIF5A_ENSG00000132507 | day1/pre-b | 0.331 | 0.0282 | 44 |
| IFN-γ/IL-5 SFC day 28 | CPPED1_ENSG00000103381 | day1/pre-b | 0.331 | 0.0282 | 44 |
| IFN-γ/IL-5 SFC pre-b | NCBP1_ENSG00000136937 | day1/pre-b | 0.32 | 0.0283 | 47 |
| IFN-γ/IL-5 SFC day 28 | DNPEP_ENSG00000123992 | day1/pre-b | 0.331 | 0.0284 | 44 |
| IFN-γ/IL-5 SFC day 28 | IFIT3_ENSG00000119917 | day1/pre-b | 0.33 | 0.0286 | 44 |
| IFN-γ/IL-5 SFC pre-b | SEC11C_ENSG00000166562 | day7/pre-b | -0.319 | 0.0287 | 47 |
| IFN-γ/IL-5 SFC day 28 | ALDOA_ENSG00000149925 | day1/pre-b | 0.33 | 0.0287 | 44 |
| IFN-γ/IL-5 SFC day 28 | NUB1_ENSG00000013374 | day1/pre-b | 0.33 | 0.0287 | 44 |
| IFN-γ/IL-5 SFC day 28 | AGTRAP_ENSG00000177674 | day1/pre-b | 0.33 | 0.0287 | 44 |
| IFN-γ/IL-5 SFC day 28 | ADRM1_ENSG00000130706 | day1/pre-b | 0.33 | 0.0289 | 44 |
| IFN-γ/IL-5 SFC pre-b | FAIM_ENSG00000158234 | day1/pre-b | -0.319 | 0.0289 | 47 |
| IFN-γ/IL-5 SFC day 28 | TM9SF4_ENSG00000101337 | day1/pre-b | 0.33 | 0.0289 | 44 |
| IFN-γ/IL-5 SFC day 28 | DUSP3_ENSG00000108861 | day1/pre-b | 0.33 | 0.0289 | 44 |
| IFN-γ/IL-5 SFC day 28 | DSE_ENSG00000111817 | day1/pre-b | 0.329 | 0.0291 | 44 |
| IFN-γ/IL-5 SFC day 28 | PLEKHM2_ENSG00000116786 | day1/pre-b | 0.329 | 0.0293 | 44 |
| IFN-γ/IL-5 SFC day 28 | X_ENSG00000276900 | day1/pre-b | 0.329 | 0.0295 | 44 |
| IFN-γ/IL-5 SFC day 28 | CTSS_ENSG00000163131 | day1/pre-b | 0.328 | 0.0295 | 44 |
| IFN-γ/IL-5 SFC day 28 | CD33_ENSG00000105383 | day1/pre-b | 0.327 | 0.03 | 44 |
| IFN-γ/IL-5 SFC pre-b | MALINC1_ENSG00000245146 | day1/pre-b | -0.316 | 0.0303 | 47 |
| IFN-γ/IL-5 SFC day 28 | BATF3_ENSG00000123685 | day1/pre-b | 0.327 | 0.0303 | 44 |
| IFN-γ/IL-5 SFC day 28 | CDKN1A_ENSG00000124762 | day1/pre-b | 0.326 | 0.0308 | 44 |
| IFN-γ/IL-5 SFC day 28 | OAS2_ENSG00000111335 | day1/pre-b | 0.325 | 0.0313 | 44 |
| IFN-γ/IL-5 SFC day 28 | GAPDH_ENSG00000111640 | day1/pre-b | 0.325 | 0.0314 | 44 |
| IFN-γ/IL-5 SFC pre-b | FXR2_ENSG00000129245 | day7/pre-b | -0.314 | 0.0314 | 47 |
| IFN-γ/IL-5 SFC day 28 | BST2_ENSG00000130303 | day1/pre-b | 0.324 | 0.0317 | 44 |
| IFN-γ/IL-5 SFC day 28 | CSF2RB_ENSG00000100368 | day1/pre-b | 0.324 | 0.0318 | 44 |
| IFN-γ/IL-5 SFC day 28 | ACP3_ENSG00000014257 | day1/pre-b | 0.324 | 0.0319 | 44 |
| IFN-γ/IL-5 SFC day 28 | LILRB2_ENSG00000131042 | day1/pre-b | 0.323 | 0.0323 | 44 |
| IFN-γ/IL-5 SFC day 28 | DDO_ENSG00000203797 | day1/pre-b | 0.323 | 0.0323 | 44 |
| IFN-γ/IL-5 SFC day 28 | MSR1_ENSG00000038945 | day1/pre-b | 0.323 | 0.0324 | 44 |
| IFN-γ/IL-5 SFC day 28 | SAP30L_ENSG00000164576 | day7/pre-b | -0.323 | 0.0324 | 44 |
| IFN-γ/IL-5 SFC pre-b | TRAV17_ENSG00000211797 | day1/pre-b | 0.312 | 0.0325 | 47 |
| IFN-γ/IL-5 SFC day 28 | RNF144B_ENSG00000137393 | day1/pre-b | 0.323 | 0.0326 | 44 |
| IFN-γ/IL-5 SFC day 28 | CLIC1_ENSG00000213719 | day1/pre-b | 0.322 | 0.0329 | 44 |
| IFN-γ/IL-5 SFC day 28 | ATP6V0D1_ENSG00000159720 | day1/pre-b | 0.322 | 0.033 | 44 |
| IFN-γ/IL-5 SFC pre-b | IFI35_ENSG00000068079 | day1/pre-b | 0.311 | 0.0332 | 47 |
| IFN-γ/IL-5 SFC day 28 | RUFY4_ENSG00000188282 | day1/pre-b | 0.322 | 0.0333 | 44 |
| IFN-γ/IL-5 SFC day 28 | PLIN3_ENSG00000105355 | day1/pre-b | 0.321 | 0.0333 | 44 |

| | | | | | |
|---|---|---|---|---|---|
| IFN-γ/IL-5 SFC pre-b | PSME2P3_ENSG00000248988 | day1/pre-b | 0.311 | 0.0335 | 47 |
| IFN-γ/IL-5 SFC day 28 | TMEM150B_ENSG00000180061 | day1/pre-b | 0.321 | 0.0335 | 44 |
| IFN-γ/IL-5 SFC day 28 | KIF2C_ENSG00000142945 | day7/pre-b | 0.321 | 0.0337 | 44 |
| IFN-γ/IL-5 SFC pre-b | IGKJ1_ENSG00000211597 | day7/pre-b | -0.31 | 0.0338 | 47 |
| IFN-γ/IL-5 SFC day 28 | LY6E_ENSG00000160932 | day1/pre-b | 0.321 | 0.0338 | 44 |
| IFN-γ/IL-5 SFC day 28 | TNFAIP2_ENSG00000185215 | day1/pre-b | 0.32 | 0.0343 | 44 |
| IFN-γ/IL-5 SFC pre-b | HLA.DRB6_ENSG00000229391 | day1/pre-b | 0.309 | 0.0347 | 47 |
| IFN-γ/IL-5 SFC day 28 | SRA1_ENSG00000213523 | day1/pre-b | 0.319 | 0.0347 | 44 |
| IFN-γ/IL-5 SFC day 28 | NAMPT_ENSG00000105835 | day1/pre-b | 0.318 | 0.0353 | 44 |
| IFN-γ/IL-5 SFC day 28 | CALCOCO2_ENSG00000136436 | day1/pre-b | 0.318 | 0.0354 | 44 |
| IFN-γ/IL-5 SFC day 28 | SIRPB1_ENSG00000101307 | day1/pre-b | 0.318 | 0.0355 | 44 |
| IFN-γ/IL-5 SFC pre-b | C2_ENSG00000166278 | day1/pre-b | 0.307 | 0.0357 | 47 |
| IFN-γ/IL-5 SFC day 28 | HCLS1_ENSG00000180353 | day1/pre-b | 0.317 | 0.0358 | 44 |
| IFN-γ/IL-5 SFC day 28 | AHSA2P_ENSG00000173209 | day7/pre-b | -0.317 | 0.0359 | 44 |
| IFN-γ/IL-5 SFC day 28 | X_ENSG00000269981 | day1/pre-b | 0.317 | 0.036 | 44 |
| IFN-γ/IL-5 SFC day 28 | G6PD_ENSG00000160211 | day1/pre-b | 0.317 | 0.036 | 44 |
| IFN-γ/IL-5 SFC day 28 | GADD45B_ENSG00000099860 | day1/pre-b | 0.317 | 0.036 | 44 |
| IFN-γ/IL-5 SFC day 28 | TTC9_ENSG00000133985 | day7/pre-b | -0.317 | 0.036 | 44 |
| IFN-γ/IL-5 SFC day 28 | FERMT3_ENSG00000149781 | day1/pre-b | 0.317 | 0.0363 | 44 |
| IFN-γ/IL-5 SFC day 28 | DDX19B_ENSG00000157349 | day1/pre-b | 0.317 | 0.0363 | 44 |
| IFN-γ/IL-5 SFC day 28 | PDLIM5_ENSG00000163110 | day1/pre-b | 0.317 | 0.0363 | 44 |
| IFN-γ/IL-5 SFC day 28 | PDCD1LG2_ENSG00000197646 | day1/pre-b | 0.317 | 0.0363 | 44 |
| IFN-γ/IL-5 SFC day 28 | SLC31A2_ENSG00000136867 | day1/pre-b | 0.316 | 0.0364 | 44 |
| IFN-γ/IL-5 SFC pre-b | HBB_ENSG00000244734 | day7/pre-b | 0.306 | 0.0366 | 47 |
| IFN-γ/IL-5 SFC pre-b | KIF11_ENSG00000138160 | day1/pre-b | 0.306 | 0.0366 | 47 |
| IFN-γ/IL-5 SFC day 28 | UCHL1_ENSG00000154277 | day7/pre-b | 0.316 | 0.0366 | 44 |
| IFN-γ/IL-5 SFC day 28 | TUBGCP2_ENSG00000130640 | day1/pre-b | 0.316 | 0.0369 | 44 |
| IFN-γ/IL-5 SFC day 28 | GNA13_ENSG00000120063 | day1/pre-b | 0.315 | 0.0374 | 44 |
| IFN-γ/IL-5 SFC pre-b | CD40_ENSG00000101017 | day1/pre-b | 0.305 | 0.0374 | 47 |
| IFN-γ/IL-5 SFC day 28 | CD38_ENSG00000004468 | day7/pre-b | 0.315 | 0.0375 | 44 |
| IFN-γ/IL-5 SFC day 28 | MOV10_ENSG00000155363 | day1/pre-b | 0.313 | 0.0383 | 44 |
| IFN-γ/IL-5 SFC pre-b | X_ENSG00000216775 | day7/pre-b | 0.303 | 0.0383 | 47 |
| IFN-γ/IL-5 SFC day 28 | LIPA_ENSG00000107798 | day1/pre-b | 0.313 | 0.0385 | 44 |
| IFN-γ/IL-5 SFC day 28 | P2RX1_ENSG00000108405 | day1/pre-b | 0.313 | 0.0387 | 44 |
| IFN-γ/IL-5 SFC pre-b | PPP1R7_ENSG00000115685 | day7/pre-b | -0.302 | 0.0388 | 47 |
| IFN-γ/IL-5 SFC day 28 | NRBP1_ENSG00000115216 | day1/pre-b | 0.312 | 0.0389 | 44 |
| IFN-γ/IL-5 SFC day 28 | OAZ2_ENSG00000180304 | day1/pre-b | 0.312 | 0.0394 | 44 |
| IFN-γ/IL-5 SFC day 28 | VPS72_ENSG00000163159 | day1/pre-b | 0.312 | 0.0395 | 44 |
| IFN-γ/IL-5 SFC day 28 | CSNK1D_ENSG00000141551 | day1/pre-b | 0.311 | 0.0396 | 44 |
| IFN-γ/IL-5 SFC day 28 | THEMIS2_ENSG00000130775 | day1/pre-b | 0.311 | 0.0397 | 44 |
| IFN-γ/IL-5 SFC day 28 | RAB24_ENSG00000169228 | day1/pre-b | 0.311 | 0.0397 | 44 |

| | | | | | |
|---|---|---|---|---|---|
| IFN-γ/IL-5 SFC pre-b | OAS2_ENSG00000111335 | day1/pre-b | 0.301 | 0.04 | 47 |
| IFN-γ/IL-5 SFC day 28 | C19orf38_ENSG00000214212 | day1/pre-b | 0.311 | 0.04 | 44 |
| IFN-γ/IL-5 SFC pre-b | PLSCR1_ENSG00000188313 | day1/pre-b | 0.3 | 0.0404 | 47 |
| IFN-γ/IL-5 SFC pre-b | SIMALR_ENSG00000226004 | day1/pre-b | 0.3 | 0.0405 | 47 |
| IFN-γ/IL-5 SFC day 28 | P2RX4_ENSG00000135124 | day1/pre-b | 0.31 | 0.0409 | 44 |
| IFN-γ/IL-5 SFC pre-b | ACVR1B_ENSG00000135503 | day1/pre-b | -0.299 | 0.0409 | 47 |
| IFN-γ/IL-5 SFC day 28 | FCAR_ENSG00000186431 | day1/pre-b | 0.309 | 0.0414 | 44 |
| IFN-γ/IL-5 SFC pre-b | MSR1_ENSG00000038945 | day1/pre-b | 0.299 | 0.0415 | 47 |
| IFN-γ/IL-5 SFC day 28 | ATP5F1B_ENSG00000110955 | day1/pre-b | 0.308 | 0.0417 | 44 |
| IFN-γ/IL-5 SFC day 28 | CHFR_ENSG00000072609 | day14/pre-b | 0.308 | 0.0417 | 44 |
| IFN-γ/IL-5 SFC day 28 | LINC02068_ENSG00000223387 | day1/pre-b | 0.308 | 0.0417 | 44 |
| IFN-γ/IL-5 SFC day 28 | PSME2P3_ENSG00000248988 | day1/pre-b | 0.308 | 0.042 | 44 |
| IFN-γ/IL-5 SFC pre-b | TIMP2_ENSG00000035862 | day1/pre-b | -0.298 | 0.0421 | 47 |
| IFN-γ/IL-5 SFC day 28 | IGHV1.3_ENSG00000211935 | day7/pre-b | 0.308 | 0.0422 | 44 |
| IFN-γ/IL-5 SFC day 28 | STEAP4_ENSG00000127954 | day1/pre-b | 0.307 | 0.0425 | 44 |
| IFN-γ/IL-5 SFC day 28 | GLB1_ENSG00000170266 | day1/pre-b | 0.306 | 0.0431 | 44 |
| IFN-γ/IL-5 SFC day 28 | HIF1A_ENSG00000100644 | day1/pre-b | 0.306 | 0.0431 | 44 |
| IFN-γ/IL-5 SFC pre-b | SH2D2A_ENSG00000027869 | day1/pre-b | 0.296 | 0.0433 | 47 |
| IFN-γ/IL-5 SFC day 28 | GPBAR1_ENSG00000179921 | day1/pre-b | 0.306 | 0.0433 | 44 |
| IFN-γ/IL-5 SFC day 28 | PSMB8_ENSG00000204264 | day1/pre-b | 0.306 | 0.0433 | 44 |
| IFN-γ/IL-5 SFC day 28 | IL27_ENSG00000197272 | day1/pre-b | 0.306 | 0.0435 | 44 |
| IFN-γ/IL-5 SFC day 28 | X_ENSG00000224579 | day1/pre-b | 0.306 | 0.0436 | 44 |
| IFN-γ/IL-5 SFC day 28 | DCTN1_ENSG00000204843 | day1/pre-b | 0.305 | 0.044 | 44 |
| IFN-γ/IL-5 SFC day 28 | CPQ_ENSG00000104324 | day1/pre-b | 0.305 | 0.0442 | 44 |
| IFN-γ/IL-5 SFC day 28 | GAA_ENSG00000171298 | day1/pre-b | 0.304 | 0.0448 | 44 |
| IFN-γ/IL-5 SFC pre-b | TRBV24.1_ENSG00000211750 | day1/pre-b | 0.294 | 0.045 | 47 |
| IFN-γ/IL-5 SFC pre-b | PIM1_ENSG00000137193 | day1/pre-b | 0.294 | 0.0451 | 47 |
| IFN-γ/IL-5 SFC day 28 | CAPZB_ENSG00000077549 | day1/pre-b | 0.304 | 0.0451 | 44 |
| IFN-γ/IL-5 SFC day 28 | MLF2_ENSG00000089693 | day1/pre-b | 0.304 | 0.0451 | 44 |
| IFN-γ/IL-5 SFC day 28 | NLRP3_ENSG00000162711 | day1/pre-b | 0.303 | 0.0456 | 44 |
| IFN-γ/IL-5 SFC day 28 | SDE2_ENSG00000143751 | day1/pre-b | 0.302 | 0.0463 | 44 |
| IFN-γ/IL-5 SFC day 28 | NANS_ENSG00000095380 | day1/pre-b | 0.302 | 0.0465 | 44 |
| IFN-γ/IL-5 SFC day 28 | TTC9_ENSG00000133985 | day1/pre-b | -0.302 | 0.0466 | 44 |
| IFN-γ/IL-5 SFC pre-b | TAP1_ENSG00000168394 | day1/pre-b | 0.291 | 0.0469 | 47 |
| IFN-γ/IL-5 SFC pre-b | TYMS_ENSG00000176890 | day7/pre-b | 0.291 | 0.0472 | 47 |
| IFN-γ/IL-5 SFC day 28 | X_ENSG00000203279 | day1/pre-b | -0.301 | 0.0472 | 44 |
| IFN-γ/IL-5 SFC day 28 | RALB_ENSG00000144118 | day1/pre-b | 0.3 | 0.0475 | 44 |
| IFN-γ/IL-5 SFC pre-b | RHOXF1P1_ENSG00000234493 | day7/pre-b | 0.29 | 0.0481 | 47 |
| IFN-γ/IL-5 SFC pre-b | DEFA3_ENSG00000239839 | day7/pre-b | 0.29 | 0.0482 | 47 |
| IFN-γ/IL-5 SFC day 28 | OASL_ENSG00000135114 | day1/pre-b | 0.299 | 0.0484 | 44 |
| IFN-γ/IL-5 SFC day 28 | SPIB_ENSG00000269404 | day1/pre-b | 0.299 | 0.0485 | 44 |

| | | | | | |
|---|---|---|---|---|---|
| IFN-γ/IL-5 SFC pre-b | X_ENSG00000271737 | day1/pre-b | -0.289 | 0.0486 | 47 |
| IFN-γ/IL-5 SFC day 28 | MGAT1_ENSG00000131446 | day1/pre-b | 0.299 | 0.0488 | 44 |
| IFN-γ/IL-5 SFC day 28 | PARP14_ENSG00000173193 | day1/pre-b | 0.299 | 0.0488 | 44 |
| IFN-γ/IL-5 SFC pre-b | MT2A_ENSG00000125148 | day1/pre-b | 0.289 | 0.0488 | 47 |
| IFN-γ/IL-5 SFC pre-b | KARS1_ENSG00000065427 | day1/pre-b | 0.289 | 0.0492 | 47 |
| IFN-γ/IL-5 SFC day 28 | REC8_ENSG00000100918 | day1/pre-b | 0.298 | 0.0494 | 44 |
| IFN-γ/IL-5 SFC day 28 | SERTAD3_ENSG00000167565 | day1/pre-b | 0.298 | 0.0496 | 44 |
| IFN-γ/IL-5 SFC day 28 | APOL3_ENSG00000128284 | day1/pre-b | 0.298 | 0.0498 | 44 |

**Supplementary Table 4**. Spearman correlation statistics between plasma cytokine changes (post/pre-b) and T cell polarization (IFN-γ/IL-5 SFC) pre- and 28 days post-booster. Significant correlates are shown (P<0.05).

| Parameter 1: Plasma cytokine changes | Parameter 2: Th1 polarization | r | *P* | n |
|---|---|---|---|---|
| IFNG (d14/pre-b) | IFN-γ/IL-5 SFC day 28 | 0.453 | 0.001 | 47 |
| IL27 (d14/pre-b) | IFN-γ/IL-5 SFC day 28 | 0.433 | 0.002 | 47 |
| CXCL9 (d14/pre-b) | IFN-γ/IL-5 SFC day 28 | 0.419 | 0.003 | 47 |
| CCL4 (d14/pre-b) | IFN-γ/IL-5 SFC day 28 | 0.388 | 0.007 | 47 |
| CXCL11 (d14/pre-b) | IFN-γ/IL-5 SFC day 28 | 0.350 | 0.016 | 47 |
| IFNG (d1/pre-b) | IFN-γ/IL-5 SFC day 28 | 0.339 | 0.021 | 46 |
| HGF (d14/pre-b) | IFN-γ/IL-5 SFC day 28 | 0.322 | 0.027 | 47 |
| FLT3LG (d14/pre-b) | IFN-γ/IL-5 SFC day 28 | 0.312 | 0.033 | 47 |
| IL2 (d3/pre-b) | IFN-γ/IL-5 SFC day 28 | 0.333 | 0.034 | 41 |
| CCL3 (d14/pre-b) | IFN-γ/IL-5 SFC day 28 | 0.303 | 0.038 | 47 |
| OLR1 (d14/pre-b) | IFN-γ/IL-5 SFC day 28 | 0.299 | 0.041 | 47 |
| CXCL11 (d3/pre-b) | IFN-γ/IL-5 SFC day 28 | 0.300 | 0.043 | 46 |
| CXCL9 (d3/pre-b) | IFN-γ/IL-5 SFC day 28 | 0.294 | 0.045 | 47 |
| IL27 (d1/pre-b) | IFN-γ/IL-5 SFC pre-b | 0.328 | 0.023 | 48 |
| IL27 (d14/pre-b) | IFN-γ/IL-5 SFC pre-b | 0.327 | 0.021 | 50 |
| CXCL9 (d3/pre-b) | IFN-γ/IL-5 SFC pre-b | 0.300 | 0.034 | 50 |
| CXCL9 (d14/pre-b) | IFN-γ/IL-5 SFC pre-b | 0.331 | 0.019 | 50 |
| CCL11 (d14/pre-b) | IFN-γ/IL-5 SFC pre-b | 0.380 | 0.006 | 50 |
| HGF (d7/pre-b) | IFN-γ/IL-5 SFC pre-b | 0.350 | 0.014 | 49 |
| CXCL10 (d14/pre-b) | IFN-γ/IL-5 SFC pre-b | 0.290 | 0.041 | 50 |
| IFNG (d1/pre-b) | IFN-γ/IL-5 SFC pre-b | 0.473 | 0.001 | 48 |
| IFNG (d3/pre-b) | IFN-γ/IL-5 SFC pre-b | 0.329 | 0.019 | 50 |
| IFNG (d7/pre-b) | IFN-γ/IL-5 SFC pre-b | 0.286 | 0.046 | 49 |
| IFNG (d14/pre-b) | IFN-γ/IL-5 SFC pre-b | 0.446 | 0.001 | 50 |
| TNF (d1/pre-b) | IFN-γ/IL-5 SFC pre-b | 0.338 | 0.019 | 48 |
| VEGFA (d14/pre-b) | IFN-γ/IL-5 SFC pre-b | 0.311 | 0.028 | 50 |
| OSM (d7/pre-b) | IFN-γ/IL-5 SFC pre-b | 0.299 | 0.037 | 49 |
| CCL4 (d7/pre-b) | IFN-γ/IL-5 SFC pre-b | 0.331 | 0.020 | 49 |
| CCL4 (d14/pre-b) | IFN-γ/IL-5 SFC pre-b | 0.320 | 0.023 | 50 |
| CXCL11 (d3/pre-b) | IFN-γ/IL-5 SFC pre-b | 0.367 | 0.010 | 49 |
| CXCL11 (d14/pre-b) | IFN-γ/IL-5 SFC pre-b | 0.334 | 0.018 | 50 |

**Supplementary Table 5**. Antibodies and dye details.

| Target | Conjugate | Host | Target | Clone | Catalog | Vendor | Application | Dilution |
|---|---|---|---|---|---|---|---|---|
| IgG | PE | Mouse | Human | JDC-10 | 9040-09 | SouthernBiotech | Ab response | 1/50 |
| IgG1 | PE | Mouse | Human | HP6001 | 9054-09 | SouthernBiotech | Ab response | 1/250 |
| IgG2 | PE | Mouse | Human | HP6025 | 9070-09 | SouthernBiotech | Ab response | 1/50 |
| IgG3 | PE | Mouse | Human | HP6050 | 9210-09 | SouthernBiotech | Ab response | 1/50 |
| IgG4 | PE | Mouse | Human | HP6025 | 9200-09 | SouthernBiotech | Ab response | 1/50 |
| CD4 | APC-eF780 | Mouse | Human | RPA-T4 | 47-0049-42 | LIFE TECH | AIM Assay | 1/50 |
| CD3 | AF700 | Mouse | Human | UCHT1 | 56-0038-42 | eBioscience | AIM Assay | 1/50 |
| CD8 | V500 | Mouse | Human | RPA-T8 | 560774 | BD | AIM Assay | 1/100 |
| CD14 | V500 | Mouse | Human | M5E2 | 561391 | BD | AIM Assay | 1/100 |
| CD19 | V500 | Mouse | Human | HIB19 | 561121 | BD | AIM Assay | 1/100 |
| CD45RA | eF450 | Mouse | Human | HI100 | 48-0458-42 | Invitrogen | AIM Assay | 1/50 |
| CCR7 | PerCP-Cy5.5 | Mouse | Human | G043H7 | 353200 | Biolegend | AIM Assay | 1/25 |
| OX40 | PE-Cy7 | Mouse | Human | Ber-ACT35 | 350012 | Biolegend | AIM Assay | 1/50 |
| CD137 | APC | Mouse | Human | 4B4-1 | 309810 | Biolegend | AIM Assay | 1/50 |
| CD25 | FITC | Mouse | Human | M-A251 | 555431 | BD | AIM Assay | 1/50 |
| CD69 | BV605 | Mouse | Human | FN50 | 562989 | BD | AIM Assay | 1/50 |
| PDL1 | PE | Mouse | Human | 29E.2A3 | 329706 | Biolegend | AIM Assay | 1/50 |
| Viability Dye | eF506 | - | All species | - | 65-0866-14 | Thermo Fisher | AIM Assay | 1/500 |
